# A practical computerized decision support system for predicting the severity of Alzheimer’s disease of an individual

**DOI:** 10.1101/573899

**Authors:** Magda Bucholc, Xuemei Ding, Haiying Wang, David H. Glass, Hui Wang, Girijesh Prasad, Liam P. Maguire, Anthony J. Bjourson, Paula L. McClean, Stephen Todd, David P. Finn, KongFatt Wong-Lin, for the Alzheimer’s Disease Neuroimaging Initiative

## Abstract

Computerized clinical decision support systems can help to provide objective, standardized, and timely dementia diagnosis. However, current computerized systems are mainly based on the group analysis, discrete classification of disease stages, or expensive and not readily accessible biomarkers, while current clinical practice relies relatively heavily on cognitive and functional assessments (CFA). In this study, we developed a computational framework using a suite of machine learning tools for identifying key markers in predicting the severity of Alzheimer’s disease (AD) from a large set of biological and clinical measures. Six machine learning approaches, namely Kernel Ridge Regression (KRR), Support Vector Regression (SVR), and k-Nearest Neighbor (kNN_reg_) for regression and Support Vector Machine (SVM), Random Forest (RF), and k-Nearest Neighbor (kNN_class_) for classification, were used for the development of predictive models. We demonstrated high predictive power of CFA. Predictive performance of models incorporating CFA was shown to be consistently higher accuracy than those based solely on biomarker modalities. We found that KRR and SVM were the best performing regression and classification methods respectively. The optimal SVM performance was observed for a set of four CFA test scores (FAQ, ADAS13, MoCA, MMSE) with multi-class classification accuracy of 83.0%, 95%CI = (72.1%, 93.8%) while the best performance of the KRR model was reported with combined CFA and MRI neuroimaging data, i.e., *R^2^* = 0.874, 95%CI = (0.827, 0.922). Given the high predictive power of CFA and their widespread use in clinical practice, we then designed a data-driven and self-adaptive computerized clinical decision support system (CDSS) prototype for evaluating the severity of AD of an individual on a continuous spectrum. The system implemented an automated computational approach for data pre-processing, modelling, and validation and used exclusively the scores of selected cognitive measures as data entries. Taken together, we have developed an objective and practical CDSS to aid AD diagnosis.

## 1. Introduction

Recent advances in machine learning (ML) and big data analytics have led to the emergence of a new generation of clinical decision support systems (CDSSs) designed to exploit the potentials of data-driven decision making in patient monitoring, particularly in the area of internal medicine, general practice, and remote monitoring of vital signs (Gálvez et al., 2013, Helldén et al., 2015 Lisboa & Taktak, 2006, Skyttberg, Vicente, Chen, Blomqvist, & Koch, 2016). Improved access to large and heterogeneous healthcare data and an integration of advanced computational procedures into CDSSs has enabled the real-time discovery of similarity metrics for patient stratification, development of predictive analytics for risk assessment, and selection of patient-specific therapeutic interventions at the time of decision-making (Brown, 2016, Dagliati et al., 2018, Farran, Channanath, Behbehani, & Thanaraj, 2013). CDSSs provide clinical decision support at the time and location of care rather than prior to or after the patient encounter and therefore, help streamline the workflow for clinicians and assist real-time decision-making (diagnosis, prognosis, treatment) (Castaneda et al., 2015, Wright et al., 2016). Numerous studies demonstrated that CDSSs contributed to improving patient safety and care by decreasing the number of therapeutic and diagnostic errors that are unavoidable in human clinical practice (Lindquist, Johansson, Petersson, Saveman, & Nilsson, 2008) and reduced the workload of medical staff, especially in contexts that require frequent monitoring or complex decision-making, such as management of chronic diseases (Wright et al., 2016). Current research directions in dementia, with Alzheimer’s disease (AD) being its most common form, focuses on interventions and treatments that can modify progression of dementia symptoms or lead to an early identification of individuals at risk of developing dementia (Brodaty et al., 2016, Ritchie et al., 2017). Increasing evidence suggests that early diagnosis of dementia can lead to significant clinical and economic benefits. However, the underdiagnosis of dementia is currently one of the key deficiencies in disease management in the primary care setting (Dodd, Cheston, & Ivanecka, 2015, Lang et al., 2017, Paterson & Pond, 2009). Research indicates that low dementia detection rates in primary care are mainly related to the absence of standardized and reliable screening tools, inadequate training on dementia of general practitioners (GPs), and the GPs’ lack of confidence in providing a correct diagnosis (Koch, Iliffe, & EVIDEM-ED project, 2010).

Technology-based tools have considerable potential to transform the dementia care pathway. CDSS utilized in the early diagnosis of AD may allow for the selection of patients for clinical trials at the earliest possible stage of disease development and enable clinicians to initiate the treatment as early in the disease process as possible to more effectively arrest or slow disease progression. A number of applications have been developed to serve as enabling tools for dementia diagnostics (Mandala, Saharana, Khana & Jamesa, 2015). These include software applications that provide practical information for those caring for dementia patients (e.g., Dementia Support by Swedish Care International, Alzheimer’s and Other Dementias Daily Companion, MindMate) as well as tools used for mobile cognitive screening (e.g., MOBI-COG, Mobile Cognitive Screening, Dementia Screener, Sea Hero Quest, CANTAB). In addition, CDSSs, designed to aid clinical decision making by adapting computerized clinical practice guidelines to individual patient characteristics or integrating machine learning methodologies for pattern recognition, have been recently gaining more interest in expediting dementia diagnosis and disease management (Antila et al., 2013, Frame, LaMantia, Bynagari, Dexter, & Boustani, 2013, Lindgren, 2011, Lindgren, Eklund, & Eriksson, 2002). It has been shown that such systems are more sensitive in detecting an early-stage disease and more objective than diagnostic decisions made by a single practitioner (Moja et al., 2015).

Despite the fact that advanced computational approaches for AD classification and progression have been applied to large sets of patient data, including magnetic resonance imaging (MRI) (Karas et al., 2008, Lebedeva et al., 2017, Moradi et al., 2015), positron emission tomography (PET) (Higdon et al., 2004, Grimmer et al., 2016, Sanchez-Catasus et al., 2017), cerebrospinal fluid (CSF) biomarkers (Forlenza et al., 2015, Handels et al., 2017, Mattsson et al., 2009), combination of the neuroimaging modalities (Youssofzadeh, McGuinness, Maguire, & Wong-Lin., 2017), and cognitive and functional assessments (CFA) (Ding et al., 2018, Chapman et al., 2011, Korolev, Symonds, Bozoki & Alzheimer’s Disease Neuroimaging Initiative, 2016, Maroco et al., 2011), there is a significant gap between research outputs and their actual utilization in daily clinical practice. In contrast to other disease areas, the integration of machine learning methodologies into CDSSs and their deployment for a routine use in AD diagnostics is still very rare. The few systems that are used in dementia diagnostics require information from expensive and labour-intensive biomarkers (Antila et al., 2013, Soininen et al., 2012) or implement predictive methodologies based on discrete classes for the different stages of the disease even if the underlying neurobiology could possibly evolve in a continuous manner (Onoda & Yamaguchi, 2014). Furthermore, to the best of our knowledge, no CDSS for dementia detection or management has been developed so far for the use in the primary care setting.

The aim of this study is two-fold: (1) to describe the developmental process of a computational framework for identifying key measures in predicting the severity of AD; and (2) to build upon this framework to develop a data-driven and self-adaptive prototype of a CDSS for evaluating the severity of AD of an individual on a continuous spectrum. In order to achieve this, we first utilize a suite of machine learning techniques to extract useful information from large volumes of patient data and provide a disease outcome prediction for different types and combinations of AD markers. We demonstrate that CFA can reliably and accurately provide prediction of AD severity. Next, we design a CDSS that incorporates an automated computational approach for data pre-processing, modelling, and validation and uses selected CFA scores as data input. Since our system was designed to utilize information from readily available and cost-effective CFA markers, it can be easily implemented in general clinical practice.

## 2. Material and methods

### 2.1 Development of a computational framework

#### 2.1.1 Participants

Patient records from the Alzheimer’s Disease Neuroimaging Initiative (ADNI) database (adni.loni.usc.edu) were used to develop the computational approach for evaluating the cognitive decline of an individual associated with AD. The primary goal of ADNI has been to test whether MRI, PET, other biological markers, and clinical and neuropsychological assessments can be combined to measure the progression of mild cognitive impairment (MCI) and early AD.

The Clinical Dementia Rating Sum of Boxes (CDRSB) scores of 488 patients with a complete dataset of structural MRI and PET imaging, CSF biomarkers, CFA scores, socio-demographic features and medical history were used to describe AD staging and acted as an outcome (response) measure in prediction models. The CDRSB score is widely accepted in the clinical setting as a reliable and objective AD assessment tool (Cedarbaum et al., 2013). In total, we identified 178 cognitively healthy controls (CDRSB = 0), 263 subjects with questionable cognitive impairment (QCI) (0.5 ≤ CDRSB ≤ 4.0), 46 patients with mild AD (4.5 ≤ CDRSB ≤ 9.0), and 1 patient with moderate AD (9.5 ≤ CDRSB ≤ 15.5). Since only one patient with moderate AD was identified, the subjects from mild and moderate AD categories were combined into one mild/moderate AD category.

#### 2.1.2 Data types

We considered 66 features as potential predictors of cognitive decline associated with AD including 38 assessments/biomarkers (10 clinical and 28 biological measures) and 28 risk factors (family history, medical history, and sociodemographic characteristics). The cognitive and functional assessments offered information on memory deficits and behavioural symptoms of AD, CSF measures corresponded to the pathological changes at the biological level, while neuroimaging features allowed us to evaluate the neural degeneration related to AD. Sociodemographic, family, and patient’s medical history data enabled the identification of risk factors associated with increased risk of developing AD.

Clinical measures included: Mini-Mental State Examination (MMSE) (Folstein, Robins & Helzer, 1983); Alzheimer’s Disease Assessment Scale 13 (ADAS13) (Mohs et al., 1997); Montreal Cognitive Assessment (MoCA) (Nasreddine et al. 2005); Logical Memory – Immediate Recall (LIMMTOTAL) (Abikoff et al., 1987); Logical Memory – Delayed Recall (LDELTOTAL) (Abikoff et al., 1987); Rey Auditory Verbal Learning Test (RAVLT): Immediate, Learning, Forgetting, and Perc Forgetting (Rey, 1964); and Functional Assessment Questionnaire (FAQ) (Pfeffer, Kurosaki, Harrah Jr, Chance & Filos, 1982).

Biological data consisted of neuroimaging measurements and CSF biomarkers. Neuroimaging measures utilized MRI and PET (FDG and 18F-AV-45) data. MRI measures included volumetric data of hippocampus, ventricles, entorhinal, fusiform gyrus, middle temporal gyrus (MidTemp), whole brain, and intracerebral volume (ICV). The regional brain volumes were normalized by ICV. We also considered the volumetric data of intracranial gray matter (GRAY), white matter (WHITE), cerebrospinal fluid (CSF_V), and white matter hyperintensities (WHITMATHYP). Furthermore, two Boundary Shift Integral (BSI) measures were evaluated: whole brain (BRAINVOL) and ventricle (VENTVOL). Finally, we analysed the Florbetapir summary data represented by the gray matter regions of interest (frontal, anterior/posterior cingulate, lateral parietal, lateral temporal) normalized by the reference region of whole cerebellum (WHOLECEREBNORM). FDG-PET (FDG) was determined as a sum of mean glucose metabolism averaged across 5 regions of interest, i.e., right and left angular gyri (Angular Right and Temporal Left respectively), bilateral posterior cingulate (CingulumPost Bilateral), right and left inferior temporal gyri (Temporal Right and Temporal Left respectively) (Landau et al., 2011). Beside the composite FDG-PET, we also considered measurements for separate FDG-ROIs (i.e., Angular Right and Left, Temporal Right and Left, CingulumPost Bilateral) (Jagust et al., 2010). 18F-AV-45 PET (AV45) was represented by the mean of Florbetapir (F-18) standardized uptake value ratios (SUVR) of frontal, anterior and posterior cingulate, lateral parietal, and lateral temporal cortex (Landau et al., 2012). Other PET measures included spatial extent of hypometabolism determined using 3-dimensional stereotactic surface projection analysis (SUMZ2, SUMZ3) (Chen et al., 2010). In addition, CSF concentrations of total tau protein - t-tau (TAU), amyloid-β peptide of 42 amino acids - Aβ_1–42_ (ABETA), and phosphorylated tau - p-tau_181p_ (PTAU) were studied, as were ratios of t-tau to Aβ_1–42_ (TAU_ABETA), and p-tau_181p_ to Aβ_1–42_ (PTAU_ABETA). The complete overview of data types used in our study and their abbreviations are shown in Table A.1.

#### 2.1.3 Feature selection and modelling approach

The development of the computational framework consists of several steps. First, we conducted feature standardization to assimilate clinical measurements of diverse scales (Liu & Motoda, 2007). Accordingly, all features were rescaled so that they had the properties of a standard normal distribution with a mean of 0 and a standard deviation of 1 (Liu & Motoda, 2007). The full dataset was then split into a model development set (90%) and a testing set (10%) was used for evaluating and comparing performances of competing models. The model development set was further split into the training and validation set. The training data was used to predict the responses for the observations in the validation set. This provided us with an unbiased evaluation of a model fit on the training dataset while tuning the hyperparameters of the model. For the validation procedure, we used the leave-one-out cross validation (LOOCV), which is a k-fold validation where *k = n* (Elisseeff & Pontil, 2003). The final model evaluation was conducted on a held out testing set that has not been used prior, either for training the model or tuning the model’s parameters (i.e., model validation).

Since machine learning algorithms tend to produce biased models when dealing with imbalanced datasets, the Synthetic Minority oversampling technique (SMOTE) was used to handle the class imbalance in the model development set by resampling original patient data and creating synthetic instances (Chawla, Bowyer, Hall & Kegelmeyer, 2002). For improved generalization performance of predictive models, feature selection was implemented to identify the most relevant subset of features for predicting AD severity. Three regression models (Kernel Ridge Regression (KRR), Support Vector Regression (SVR), and k-Nearest Neighbor Regression (kNN_reg_)) and three classification models (Support Vector Machine (SVM), Random Forest (RF), k-Nearest Neighbor Classification (kNN_class_)) were developed and their performance tested for different modality types and their combinations. The selection of features that achieved high predictive accuracy for the best performing classification and regression model was later used as entry input for CDSS. A leave-one-out cross validation (LOOCV) was applied for hyper-parameters optimization. The overall procedure for model development and evaluation is shown in Fig. 1.

**Fig. 1.**
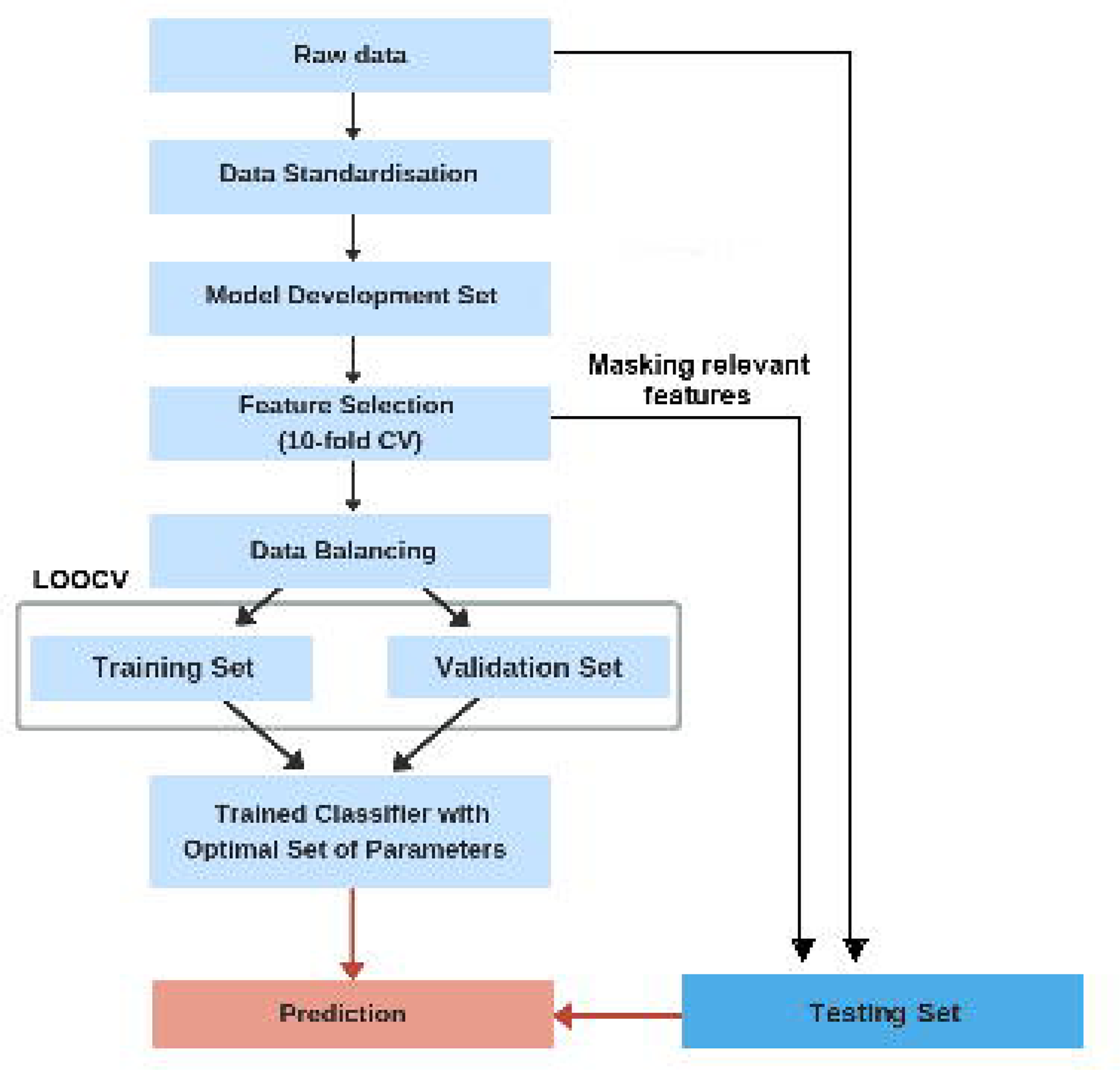
Overview of the model development and validation procedure.

##### 2.1.3.1 Feature selection

Previous studies typically used the univariate filtering methods to filter out the least promising features before the development of a predictive model (Michalak & Kwaśnicka, 2006). However, such filtering approaches can prompt loss of relevant features that are meaningless by themselves but when considered together, can improve model performance (Perez-Riverol, Kuhn, Vizcaíno, Hitz & Audain, 2017). To overcome this, the wrapper methods can be applied to assess the importance of specific feature sets. It has been shown that wrappers obtain subsets with better performance than filters. Wrappers use a search procedure to generate and evaluate different subsets of features in the space of possible feature subsets by training and testing a specific classification model (Hira & Gillies, 2015).

The commonly used classification algorithms for identifying the most relevant input variables are: Naïve Bayes (Cortizo & Giraldez, 2006, Panthong & Srivihok, 2015), SVM (Maldonado & Weber, 2009, Maldonado, Weber & Famili, 2009), Random Forest (Rodin et al., 2009), Bagging (Panthong & Srivihok, 2015), AdaBoost (Panthong & Srivihok, 2015), and Extreme Learning Machines (Benoít, Van Heeswijk, Miche, Verleysen & Lendasse, 2013). These classification techniques combined with a greedy search algorithm allow for finding the optimal number of features by iteratively selecting features based on the classifier performance (Bengio et al., 2003).

Since ADNI dataset is characterized by high dimensionality that increases the complexity of computation and analysis, we used the feature selection technique that was found to minimize redundancy and allowed for identifying features with the highest relevance to the disease class (Granitto, Furlanello, Biasioli & Gasperi, 2006). As such, we applied the Recursive Feature Elimination (RFE) method coupled with Random Forest for measuring variable importance. The RFE technique has been widely applied in healthcare applications due to its efficiency in reducing the complexity (Li, Xie, & Liu, 2018). Furthermore, studies demonstrated that RF-RFE outperformed SVM-RFE in finding small subsets of features with a high discrimination capability and required no parameter tuning to produce competitive results (Granitto, Furlanello, Biasioli & Gasperi, 2006).The RFE method with the 10-fold validation was applied on the model development set (Bengio et al., 2003). For better replicability, the 10-fold CV procedure was repeated 10 times with different partitions of the data to avoid any bias introduced by randomly partitioning dataset in the cross-validation. The RFE technique searched for the optimal combination of predictors (among all possible subsets) that maximized model performance through backward feature elimination based on the predictor importance measure as a ranking criterion. At each iteration, the Random Forest (RF) algorithm, incorporating a hierarchical decision tree structure was used to explore all possible subsets of the features and measure their importance with respect to the classification outcome (Gregorutti, Michel & Saint-Pierre, 2017). To assess the robustness of RF-RFE process in selecting optimal subset of features, we applied the RFE technique to another type of ensemble methods, namely, bootstrap aggregated (bagged) trees (BT) and compared the results (Panthong & Srivihok, 2015). Similarly, the BT-RFE performance was evaluated in a 10-fold cross-validation repeated five times with different split positions.

##### 2.1.3.2 Development of predictive models

A number of ML techniques have been used for AD detection. Classification approaches have been derived using Random Forest (RF) (Gray et al., 2013, Sarica, Cerasa & Quattrone, 2017), Logistic Regression (Barnes et al., 2010, Bauer, Cabral & Killiany, 2018, Chary et al. 2013, Wolfsgruber et al., 2014), and SVM (Casanova, Hsu, & Espeland, 2015, Cui et al., 2011, Klöppel et al., 2008, Ritter et al., 2015, Weygandt et al., 2011). In particular, the SVM showed great promise in improving diagnosis and prognosis in AD, especially in the studies characterized by a relatively small number of participants and disparate and high-dimensional data types (Dyrba, Grothe, Kirste, & Teipel, 2015, Klöppel et al., 2008, Long, Chen, Jiang, Zhang, & Alzheimer’s Disease Neuroimaging Initiative, 2017, Magnin et al., 2009). Furthermore, the SVM often outperformed other machine learning algorithms used for AD classification (e.g. RF, logistic regression) (Samper-González et al., 2018, Tripoliti, Fotiadis, Argyropoulou, & Manis, 2010).

Compared to the ML classification methods, regression approaches focus on the estimation of continuous clinical variables along the continuum of disease severity (Wang, Fan, Bhatt & Davatzikos, 2010). Several regression methods have been applied in AD studies (Duchesne, Caroli, Geroldi, Collins, & Frisoni, G. 2009, Duchesne, Caroli, Geroldi, Frisoni, & Collins, 2005, Youssofzadeh et al., 2017). However, linear regression models have been often ineffective in capturing nonlinear relationships between biomarkers (e.g. neuroimaging data) and cognitive scores, especially when limited training examples of high dimensionality were used (Duchesne et al., 2009). On the other hand, nonparametric kernel regression methods yielded relatively robust estimations of continuous variables with good generalization ability (Liu, Cao, Yang, & Zhao, 2018, Wang et al., 2010). Regularized regression techniques, such as Ridge Regression, performed especially well given high dimensional and colinear AD data (Teipel et al., 2017, Youssofzadeh et al., 2017). In addition, the Ridge Regression combined with the kernel trick demonstrated high predictive performance when applied to individual patient data (Youssofzadeh et al., 2017).

Our study built upon earlier findings and used six different non-parametric methods for the development of predictive models, namely SVM, RF, and kNN_class_ for classification and KRR, SVR, and kNN_reg_ for regression. For each selected technique, we tested a series of values for the tuning process with the optimal parameters determined based on the model performance. The results of the best performing regression and classification algorithms are presented in the main text; the results of the remaining methods can be found in the Supplementary Material (Supplementary Table A.2., A.3, and A.4).

The distinction between regression and classification models was reflected in definition of the response variable (CDRSB). The regression models predicted a numerical value from a range of continuous values (i.e., 0 < CDRSB < 15.5) while the classification models predicted the target class, i.e., ‘Normal’ (CDRSB = 0), ‘QCI’ (0.5 ≤ CDRSB ≤ 4.0), ‘Mild/Moderate’ (4.5 ≤ CDRSB ≤ 15.5). Since the model performance greatly depends on the choice of a kernel function (Hainmueller & Hazlett, 2014, Matheny, Resnic, Arora & Ohno-Machado, 2007), we tested different types of kernels, i.e., linear, polynomial, and radial basis function, and selected the one that maximized the performance measure for each model type.

###### 2.1.3.2.1 Kernel Ridge Regression

The KRR combines ridge regression with a kernel trick allowing for mapping the input space into a higher dimensional space of nonlinear functions of predictors (Murphy, 2014). The general form of the KRR is described by:

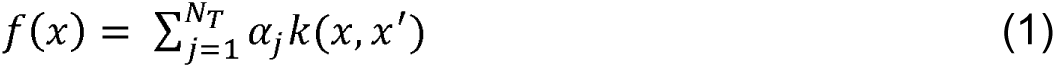

where *N_T_* is the number of training points, *k* is the kernel function, and *α* are the weights obtained through the minimization of the cost function:

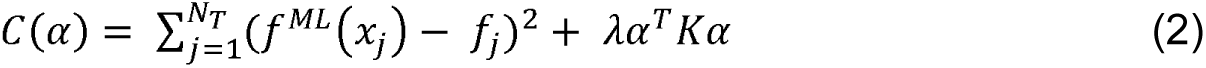

where *α* = (*α*_1_, …*α_N_T__*)*^T^*, *K* is the kernel matrix, and, *λ* controls the amount of regularization applied to the model (Vu et al., 2015). The best performance of the KRR model was achieved by applying a radial basis function (RBF) kernel defined as:

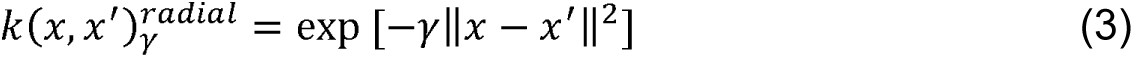

where *x* and *x′* are input vectors, and *γ* > 0 is a width parameter (Murphy, 2014).

###### 2.1.3.2.2 Support Vector Machine and Support Vector Regression

SVM is a classification technique that performs classification tasks by mapping the input vectors onto a higher dimensional space denoted as Φ: *R_d_* → *H_f_* (*d* < *f*) where an optimal separating hyperplane is constructed using a kernel function *k*(*x_i_*, *x_j_*) (Ramírez et al., 2013).

The performance of the SVM classifier was maximized using a polynomial kernel:

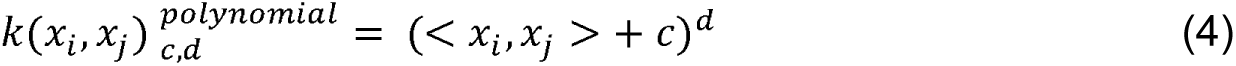

where *x_i_* and *x_j_* are vectors in the input space, *c* is a free parameter trading off the influence of higher-order versus lower-order terms in the polynomial, and *d* is the degree of polynomial (Cortes & Vapnik, 1995).

SVR is based on the same principles as SVM. In contrast to traditional regression techniques, SVR focuses on minimizing the bound of the generalization error instead of seeking to minimize the prediction error on the training set (training error) (Basak, Pal, & Patranabis, 2007). The objective of SVR is to find a regression function, *y* = *f*(*x*), such as it predicts the outputs {y} corresponding to a new input-output set {(x,y)} which are drawn from the same underlying joint probability distribution as the training set g = {(*x*_1_, *y*_1_),(*x*_2_, *y*_2_),(*x*_p_, *y*_p_)}, where *x_i_ ϵ υ^N^* is the vector of input variables and *y_i_ ϵ υ* is the vector of corresponding output values (Awad & Khanna, 2015). The basic concept of SVR is to non-linearly transform the original input space into a higher dimensional feature space and perform linear regression in this feature space by ε-insensitive loss (Awad & Khanna, 2015). The SVR ε-insensitive loss function penalizes misestimates that are farther than ε from the desired output. The ε parameter determines the width of the ε-insensitive region (tube) around the function; a lower tolerance for error is reflected in a smaller ε value. If the predicted value is within the ε-zone, the loss is zero. If the predicted value is located outside the ε-zone, the loss is defined by the magnitude of the difference between the predicted value and the ε radius (Awad & Khanna, 2015).

###### 2.1.3.2.3 k-Nearest Neighbors

kNN is a non-parametric approach applied to both classification and regression problems. The prediction of values of any new data points uses the ‘feature similarity’ measure (Kramer, 2013). Accordingly, given a predefined threshold for the rule (i.e. the k number of neighbors) a new point is assigned a value based on its distance to training examples. Here, the distance between two data points is determined using the normalized Euclidean distance function defined as:

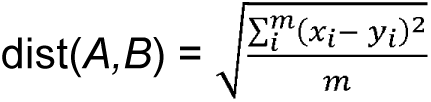

where *A* and *B* are represented by feature vectors *A* = (*x_1_*, *x_2_*, …, *x_m_*), *B* = (*y_1_, y_2_*, *…,y_m_*), and *m* is the dimensionality of the feature space (Kramer, 2013). The kNN classification assigns a class label of the majority of the k-nearest patterns in the feature space while the kNN regression calculates the mean of the function values of its k-nearest neighbors (Kramer, 2013).

###### 2.1.3.2.4 Random Forest

RF estimates the importance of features included in a model by constructing an ensemble of decision trees (Rodin et al., 2009). As a boosting type of algorithm, RF combines the efforts of an ensemble of weak classifiers to build a single, stronger classifier. It achieves it by training a specified number of decision trees using different partitions of the training set and conducting the following randomizing operations: 1) each tree is trained on a random bootstrap subset of the training data; 2) each node of a tree only uses a randomly selected subset of features. The trained decision trees then produce a single prediction by averaging the individual estimates from random subsamples of the data. More detail about the theory and mechanisms of RF is given in Breiman (2001).

##### 2.1.3.3 Model performance evaluation

The optimal subset of features identified during the feature selection process was subsequently used for training the selected regression and classification models. Both types of models were developed using 90% of the original data. The values of hyper-parameters used in constructing the models were optimized by applying grid search with LOOCV on the training data (Elisseeff & Pontil, 2003). The LOOCV technique is *N*-fold cross-validation, where *N* is the number of instances in the dataset. Although LOOCV is computationally intensive, choosing the number of folds equal to *N* gives more accurate assessment as the true size of the training set is closely mimicked and hence, the model bias is minimized (Elisseeff & Pontil, 2003). Accordingly, we tested each single held out patient record (validation set) on the classifier trained on the remaining (*N* - 1) patient observations. Note that, the optimal values of the parameters were determined separately for each model type and each modality type or their combinations (i.e., CFA, MRI, PET, CSF, Age). The predictive performance of trained models was later evaluated on an (unseen) test set randomly partitioned from the original data (10% of the original data). The test was performed once for each model constructed using different modality types and their combinations. This allowed us to identify a subset of features that was later used as entry input for the CDSS.

Two established measures for assessing the performance of regression models were used: the adjusted coefficient of determination (*R^2^*) and the Root Mean Square Error (RMSE) (Allen, 1997). For classification models, we calculated four metrics: multi-class classification accuracy (MCA), sensitivity, specificity, and area under the ROC curve (AUC) (Hand & Till, 2001). Since simple form of AUC is only used as a binary classification measure, we extended the definition of AUC to the case of multi-class problem by averaging pairwise comparisons (Hand & Till, 2001).

### 2.2 Development of clinical decision support system

The development of the computational framework described above allowed us to identify a subset of features with high discriminative power in evaluating levels of cognitive impairment in AD. These features were used as CDSS inputs for assessing AD severity of an individual (Bucholc et al. 2017, Bucholc et al., 2018). The CDSS workflow characteristics are shown in Fig. 2. The elements of the framework responsible for data pre-processing, modelling, and validation were automated and realized in the CDSS. The software prototype was developed using R version 3.4.1 and Shiny version 1.0.5. A team of domain experts including computer scientists and clinical experts was involved in the design process. To maximize system effectiveness, clarity, and guarantee efficient interaction with clinical staff, the visual representations of clinical data were displayed in concise formats that did not lower cognitive effort required to interpret them in a timely manner. Consultations with medical personnel enabled an understanding of the local context in which the system will be implemented. Furthermore, all involved parties became familiar with the rationale and methodological approach behind the development of our decision support tool. This closed-loop process between the computer scientist and clinicians helped us identify leading obstacles to the system’s adoption and routine use in clinical practice.

**Fig. 2.**
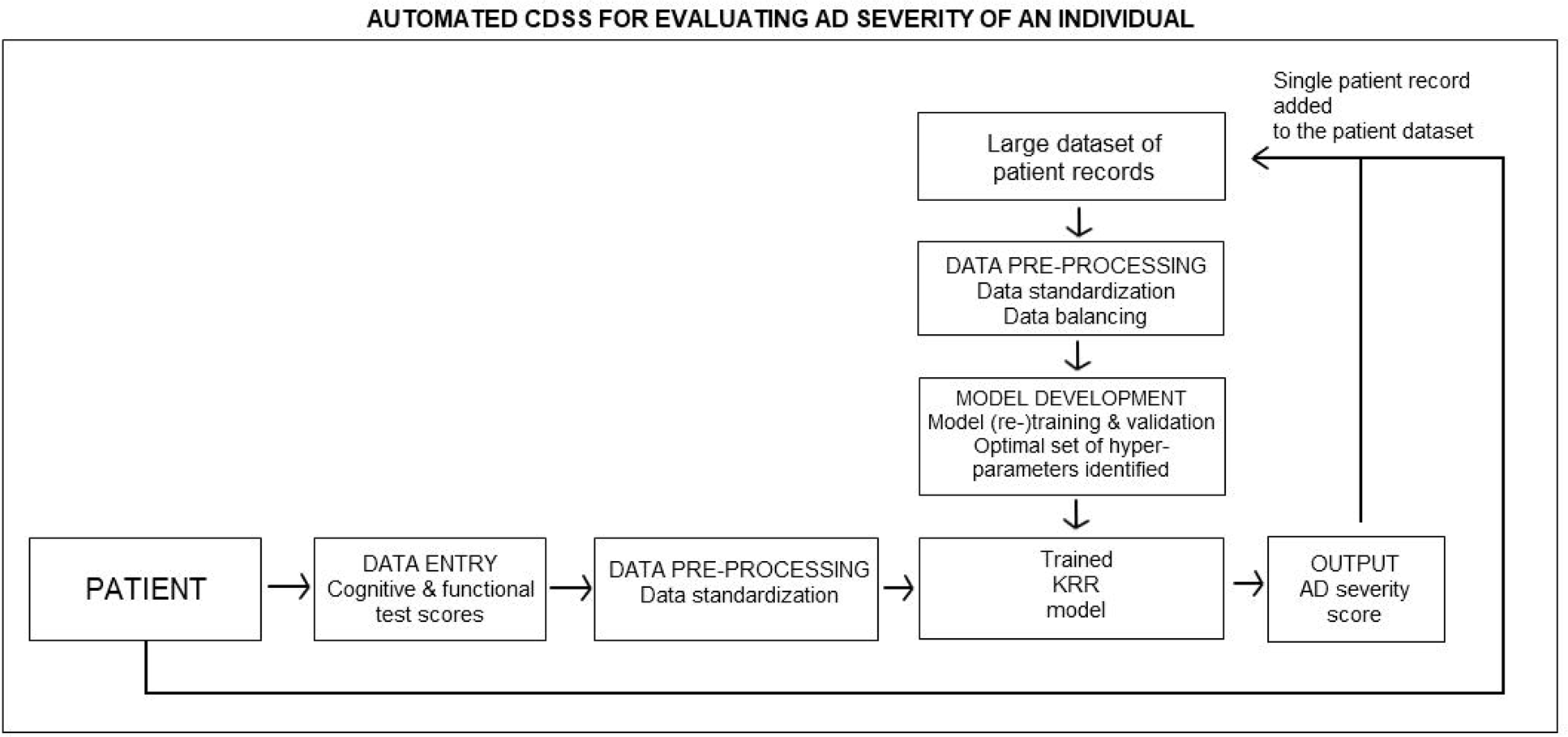
UML activity diagram of the computer-based clinical decision support system for predicting AD severity of an individual.

## 3. Results

### 3.1 Identification of AD features for the CDSS data entry

#### 3.1.1 Dimensionality reduction of AD data

Both feature selection methods (RFE-RF and RFE-BT) we consider are variants of the recursive stepwise selection approach. Fig. 3 shows the performance profile across different subset sizes evaluated with the RFE-RF (Fig. 3A) and RFE-BT (Fig. 3B) technique. The plotted values refer to the average accuracy measured using 10 repeats of 10-fold cross-validation. The accuracy of classifiers (RF and BT) was calculated for different combinations of features and the subset of features with best performance was retained.

**Fig. 3.**
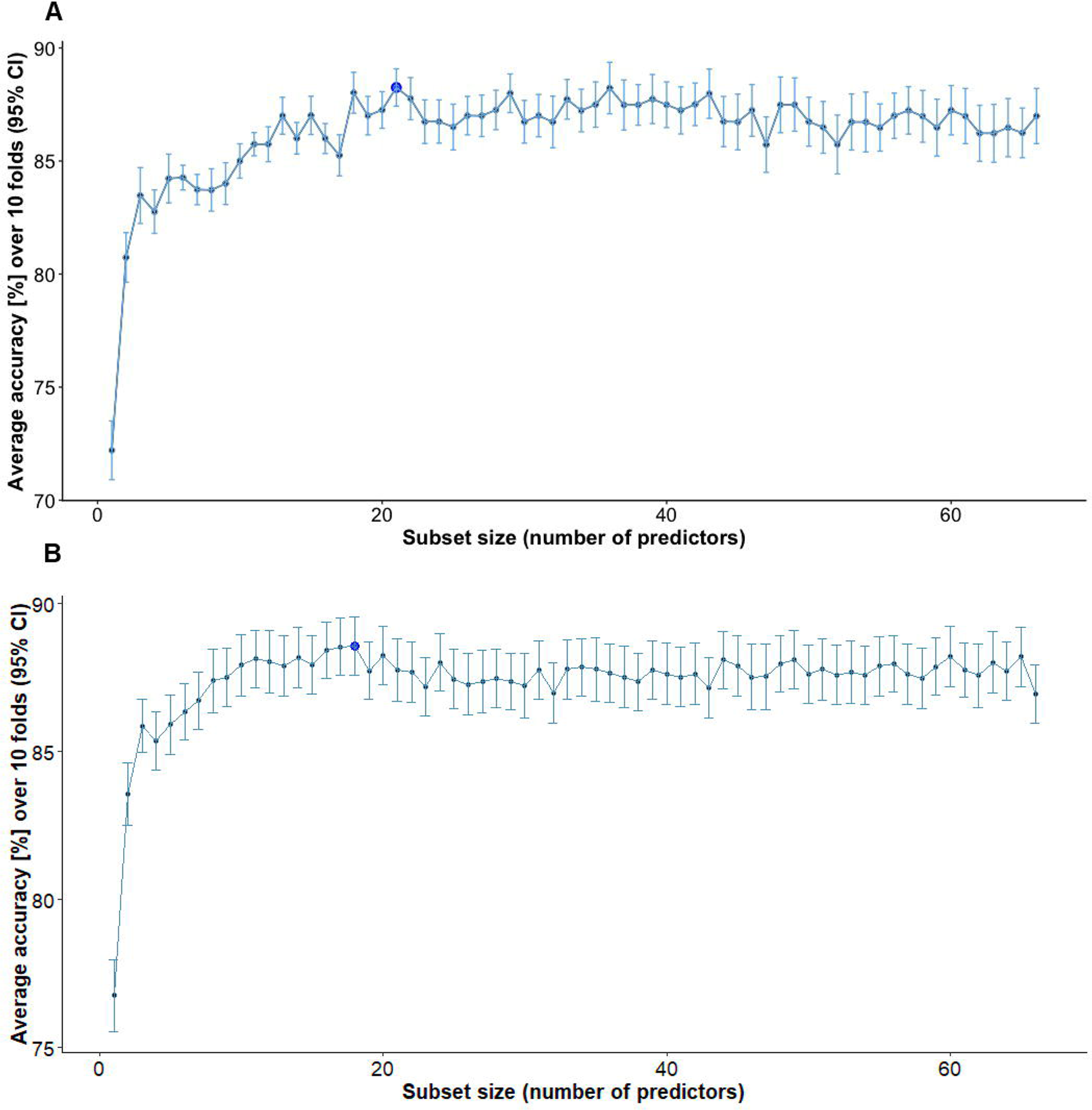
A) Performance profile across different subset sizes evaluated using the RFE-RF technique. Dark blue dot: the subset of features with the best performance B) Resampling performance of the best subset of features across different folds.

Given the RFE-RF, we found a combination of 21 features (LDELTOTAL, FAQ, MOCA, ADAS13, LIMMTOTAL, RAVLT Immediate, MMSE, Hippocampus, FDG, Angular Left, Whole Brain, Age, RAVLT Perc Forgetting, MidTemp, Angular Right, Temporal Left, SUMZ3, RAVLT Learning, TAU_ABETA, TAU, Entorhinal) to achieve the highest predictive accuracy (MCA = 88.9%, 95%CI = (88.2%, 89.6%)). The optimal subset of features identified with RFE-BT consisted of 18 features with MCA = 88.5%, 95%CI = (87.5%, 89.5%). All features (with exception of SUMZ2) selected during the RFE-BT process were also identified with RFE-RF. Since the best subset of features determined using the RFE-RF approach was more comprehensive and yielded higher accuracy, we used it for training regression and classification models.

The features identified with RF-RFE were grouped into five modality types: 1) CFA (LDELTOTAL, FAQ, MOCA, ADAS13, LIMMTOTAL, MMSE, RAVLT Immediate, RAVLT Perc Forgetting, RAVLT Learning); 2) MRI (Hippocampus, MidTemp, Entorhinal, Whole Brain); 3) PET (FDG, Angular Left, Angular Right, Temporal Left, SUMZ3); 4) CSF (TAU_ABETA, TAU); and 5) Age. The reason for grouping the features into modality types was to determine if cost-effective and non-invasive AD markers, and therefore, easier to implement into the CDSS, have high discriminative power in assessing the severity of AD. Accordingly, we analysed the performance of predictive models constructed using each data type (as well as their combinations).

#### 3.1.2 Model performance

To test the robustness of our hypothesis, we used six different ML methods for the development of predictive models, namely KRR, SVR, and kNN_reg_ for regression and SVM, RF, and kNN_class_ for classification. Our analysis showed that all models incorporating CFA into their design performed better than models based on a single or combination of biomarkers. The results of the best performing regression and classification models (KRR and SVM respectively) were presented in the main text while the performance measures for the remaining 4 models were included in the Supplementary material (Table A.2, A.3, A.4).

##### 3.1.2.1 Kernel Ridge Regression model

The KRR model constructed for a combination of CFA and biomarkers performed consistently better than models incorporating only biomarkers (either a single modality type or their combinations) (Table 1). The best performance of the KRR model was observed for the combined CFA and MRI data, i.e., *R^2^* = 0.874, 95%CI = (0.827, 0.922) (Table 1, bold). Of the two modalities, CFA features were the most discriminative while MRI markers provided complementary information about AD severity, enhancing the predictive performance of the model. Taken together, CFA provided insight into the memory deficits and behavioural symptoms of AD while MRI features offered complementary information regarding the structural degeneration of AD. Biomarker features achieved significantly lower performance, e.g., combined PET, MRI, and CSF data yielded *R^2^* = 0.417, 95%CI = (0.256, 0.578) while for PET and MRI features, we reported *R^2^*of 0.407, 95%CI = (0.237, 0.578). Given a single modality type, the model based on CFA (*R^2^* = 0.866, 95%CI = (0.809, 0.922)) clearly outperformed models constructed with MRI (*R^2^* = 0.317, 95%CI = (0.120, 0.513)), PET (*R^2^* = 0.404, 95%CI = (0.215, 0.593)) and CSF (*R^2^* = 0.024, 95%CI = (0, 0.105)) features. Modelsbuilt using Age or CSF data alone achieved the worst performance. KRR predictions of AD severity of individual patients along with the expected diagnosis for each modality type are shown in Fig. 4.

**Fig. 4.**
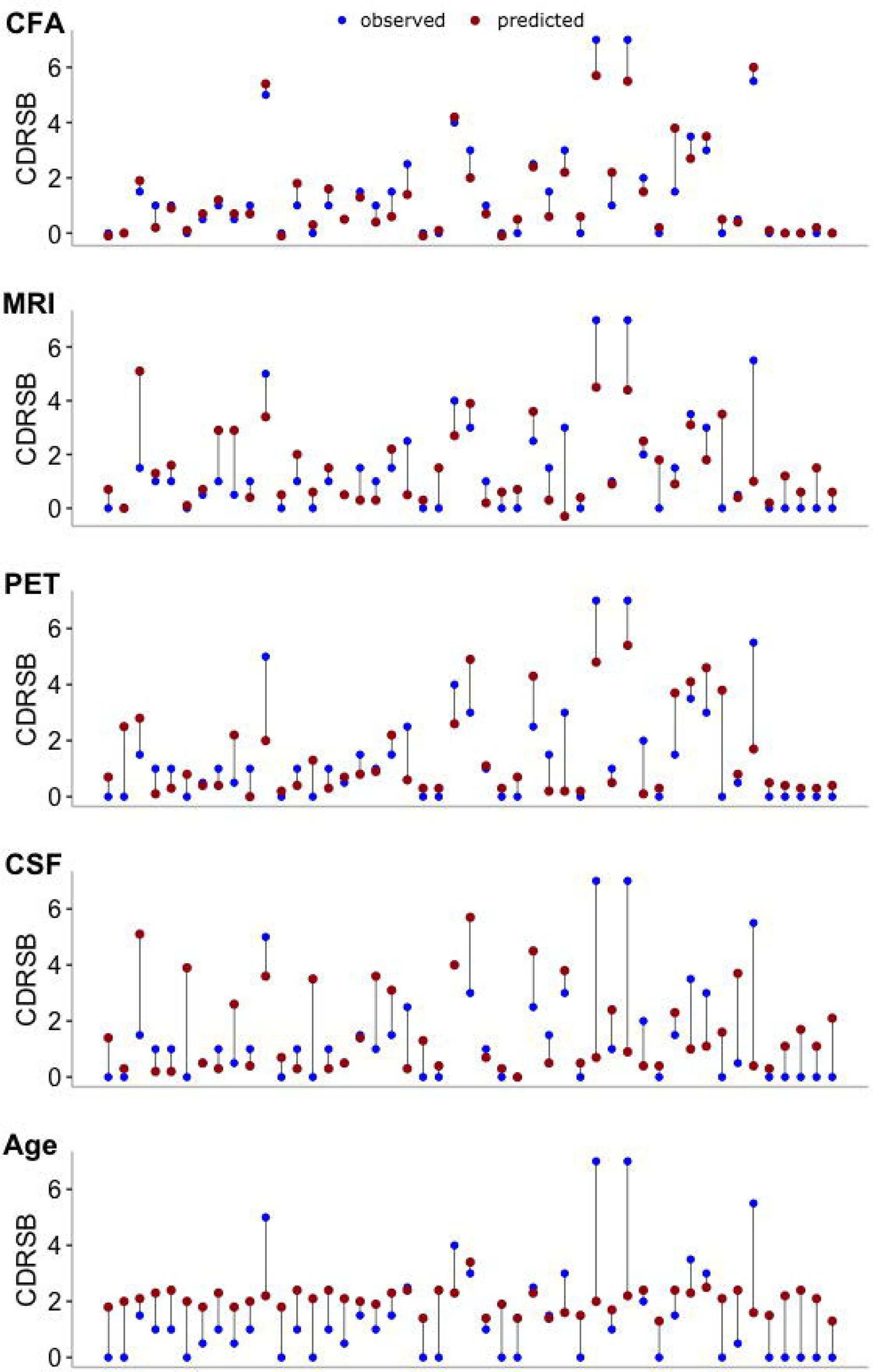
KRR model predictions of medical diagnosis (CDRSB) of individual patients for 5 modality types: a) CFA, b) MRI, c) PET, d) CSF, and e) Age. Blue dots: observed values of CDRSB; red dots: predicted values of CDRSB; vertical lines: differences between observed and predicted values of the outcome. Models’ predictions for each set of considered markers were obtained using an (unseen) testing set partitioned from the original data (10%). CFA: functional and cognitive assessments; MRI: magnetic resonance imaging; PET: positron emission tomography; CSF: cerebrospinal fluid biomarkers.

**Table 1.**
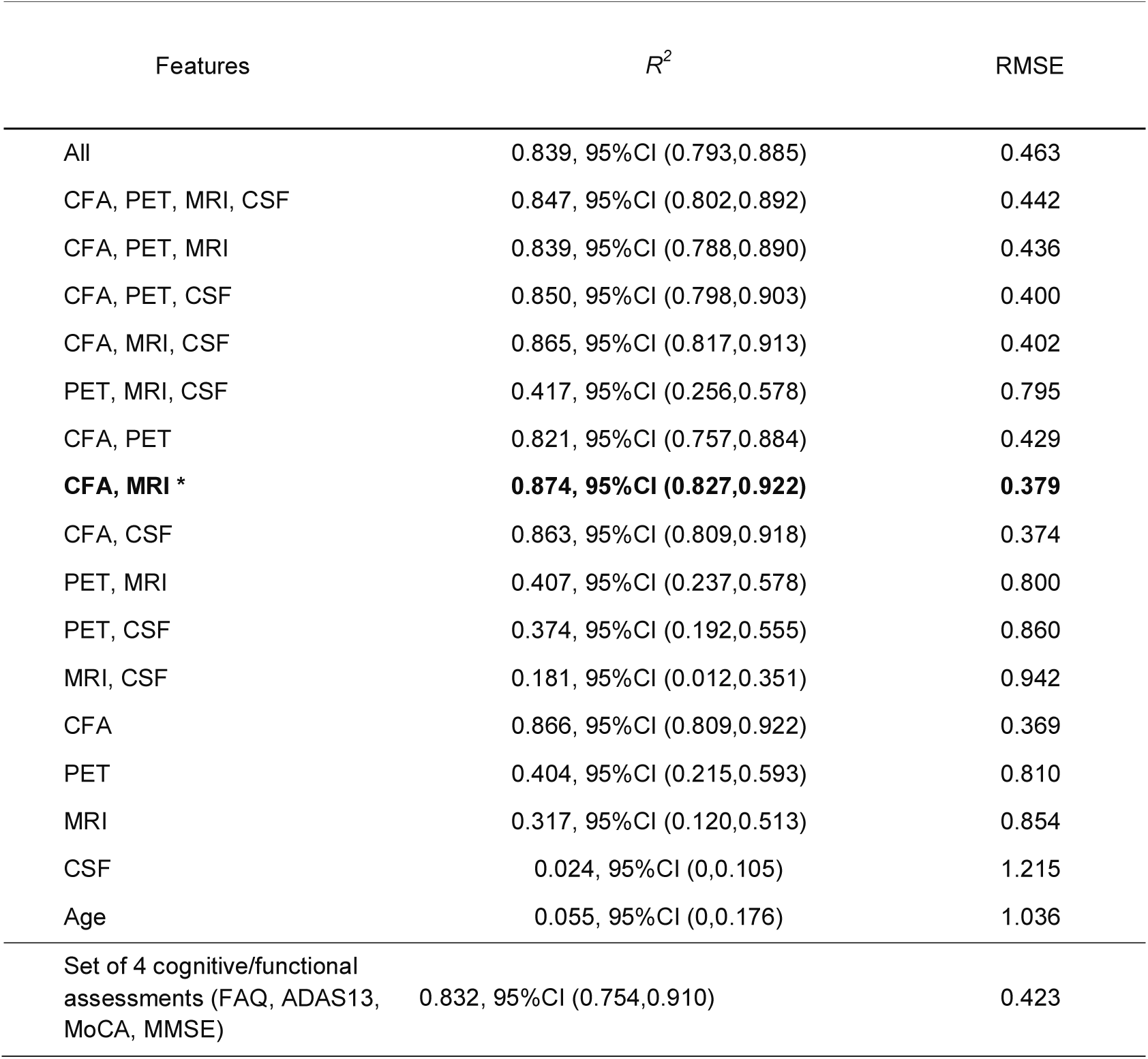
KRR model performance measures for MRI, PET, CSF and cognitive function modalities retained for the training after feature selection. CFA represents 9 selected cognitive and functional assessments (LDELTOTAL, FAQ, MOCA, ADAS13, LIMMTOTAL, MMSE, RAVLT Immediate, RAVLT Perc Forgetting, RAVLT Learning), MRI - 4 features (Hippocampus, MidTemp, Entorhinal, Whole Brain), PET – 5 features (FDG, Angular Left, Angular Right, Temporal Left, SUMZ3), and CSF – 2 features (TAU_ABETA, TAU). ‘All’ features refer to a combination of MRI, PET, CSF, CFA, and Age. Performances of predictive models for each combination of modalities were recorded using an (unseen) testing set partitioned from the original data (10% of the original data). *R^2^*: adjusted coefficient of determination; RMSE: Root Mean Square Error. Asterix (*): a subset of features with the highest *R^2^*. For more details on data types and their abbreviations, refer to Table A.1.

##### 3.1.2.2 Support Vector Machine

Three target disease classes associated with AD severity were used in SVM classification: ‘Normal’ (CDRSB = 0), ‘QCI’ (0.5 ≤ CDRSB ≤ 4.0), and ‘AD Mild/Moderate’ (4.5 ≤ CDRSB ≤ 15.5). The SVM MCA and multiclass AUC observed for a combination of all 5 modality types was 74.5%, 95%CI = (61.9%, 87.1%) and 91.6% respectively (Table 2). Again, combinations of features incorporating CFA yielded higher performance than models constructed using a single or combined biomarker modalities. The best SVM performance was observed for a subset of 4 CFA features (FAQ, ADAS13, MoCA, MMSE), i.e., MCA of 83.0%, 95%CI = (72.1%, 93.8%) and AUC = 94.9% (Table 2, bold). Given individual modality types, the model built using CFA outperformed models constructed with MRI, PET, or CSF data. Fig. 5 shows the expected diagnosis along with the corresponding SVM predictions obtained for 5 considered modality types. The best sensitivity and specificity in distinguishing Normal from QCI and Mild/Moderate AD cases was achieved for a combination of four CFA (FAQ, ADAS13, MoCA, MMSE) (sensitivity = 100% and specificity = 100%) (Table 2). The best sensitivity and specificity in identifying QCI from Normal and Mild/Moderate AD subjects was observed for combined CFA, PET, and MRI features (sensitivity = 80.8% and specificity = 85.7%). For all modality types (and their combinations), the QCI category had generally lower sensitivity than Normal and Mild/Moderate AD.

**Fig. 5.**
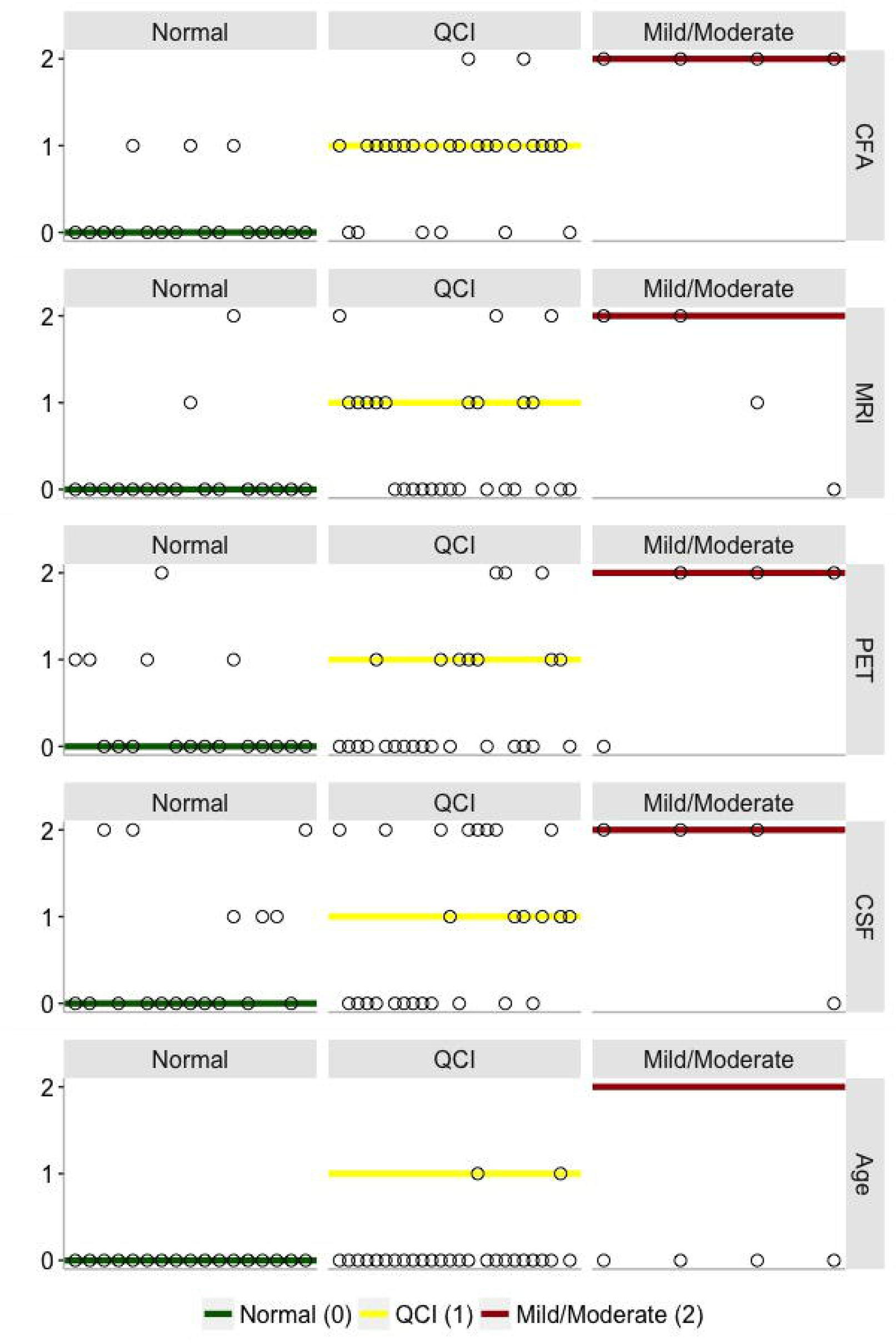
SVM model predictions of medical diagnosis of individual patients for 5 data types: a) CFA, b) MRI, c) PET, d) CSF, and e) Age. The vertical axis values and corresponding horizontal lines refer to the target CDRSB class, i.e., ‘Normal’ (green) = 0 (CDRSB = 0), ‘QCI’ (yellow) = 1 (0.5 ≤ CDRSB ≤ 4.0), and ‘Mild/Moderate’ (red) = 2 (4.5 ≤ CDRSB ≤ 15.5). Circles: predicted CDRSB class. CFA: functional and cognitive assessments; MRI: magnetic resonance imaging; PET: positron emission tomography; CSF: cerebrospinal fluid biomarkers.

**Table 2.**
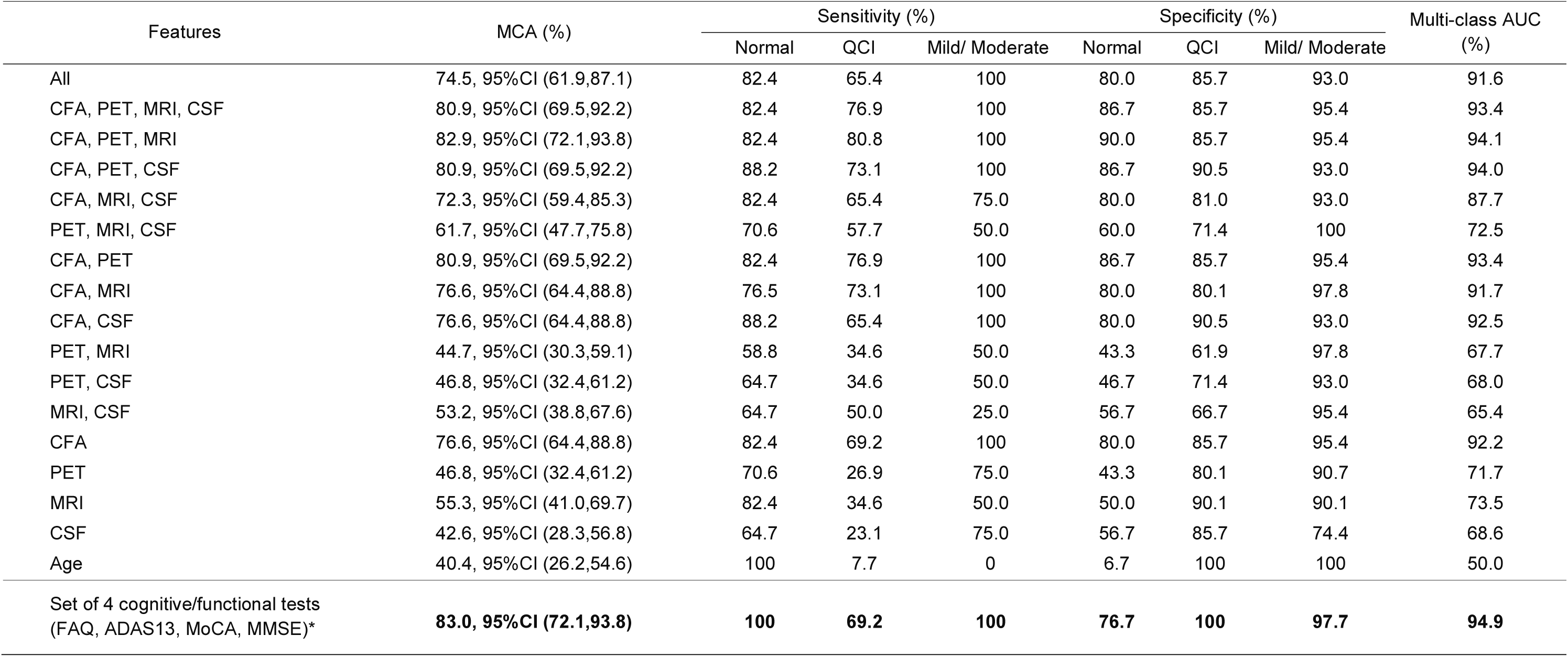
SVM model performance measures for MRI, PET, CSF and cognitive 469 function modalities retained for the training after feature selection. CFA represents 9 selected cognitive and functional assessments (LDELTOTAL, FAQ, MOCA, ADAS13, LIMMTOTAL, RAVLT Immediate, MMSE, RAVLT Perc Forgetting, RAVLT Learning), MRI – 4 features (Hippocampus, MidTemp, Entorhinal, Whole Brain), PET – 5 features (FDG, Angular Left, Angular Right, Temporal Left, SUMZ3), and CSF – 2 features (TAU_ABETA, TAU). ‘All’ features refer to a combination of MRI, PET, CSF, CFA, and Age. Performances of predictive models for each combination of modalities were recorded using an (unseen) testing set partitioned from the original data (10% of the original data). MCA: multi-class classification accuracy. Multi-class AUC: multiclass area under the curve. Asterix (*): a subset of features with the best predictive performance. For more details on data types and their abbreviations, refer to Table A.1.

| Features | MCA (%) | Sensitivity (%) |  |  | Specificity (%) |  |  | Multi-class AUC (%) |
| --- | --- | --- | --- | --- | --- | --- | --- | --- |
|  |  | Normal | QCI | Mild/ Moderate | Normal | QCI | Mild/ Moderate |  |
| All | 74.5, 95%CI (61.9,87.1) | 82.4 | 65.4 | 100 | 80.0 | 85.7 | 93.0 | 91.6 |
| CFA, PET, MRI, CSF | 80.9, 95%CI (69.5,92.2) | 82.4 | 76.9 | 100 | 86.7 | 85.7 | 95.4 | 93.4 |
| CFA, PET, MRI | 82.9, 95%CI (72.1,93.8) | 82.4 | 80.8 | 100 | 90.0 | 85.7 | 95.4 | 94.1 |
| CFA, PET, CSF | 80.9, 95%CI (69.5,92.2) | 88.2 | 73.1 | 100 | 86.7 | 90.5 | 93.0 | 94.0 |
| CFA, MRI, CSF | 72.3, 95%CI (59.4,85.3) | 82.4 | 65.4 | 75.0 | 80.0 | 81.0 | 93.0 | 87.7 |
| PET, MRI, CSF | 61.7, 95%CI (47.7,75.8) | 70.6 | 57.7 | 50.0 | 60.0 | 71.4 | 100 | 72.5 |
| CFA, PET | 80.9, 95%CI (69.5,92.2) | 82.4 | 76.9 | 100 | 86.7 | 85.7 | 95.4 | 93.4 |
| CFA, MRI | 76.6, 95%CI (64.4,88.8) | 76.5 | 73.1 | 100 | 80.0 | 80.1 | 97.8 | 91.7 |
| CFA, CSF | 76.6, 95%CI (64.4,88.8) | 88.2 | 65.4 | 100 | 80.0 | 90.5 | 93.0 | 92.5 |
| PET, MRI | 44.7, 95%CI (30.3,59.1) | 58.8 | 34.6 | 50.0 | 43.3 | 61.9 | 97.8 | 67.7 |
| PET, CSF | 46.8, 95%CI (32.4,61.2) | 64.7 | 34.6 | 50.0 | 46.7 | 71.4 | 93.0 | 68.0 |
| MRI, CSF | 53.2, 95%CI (38.8,67.6) | 64.7 | 50.0 | 25.0 | 56.7 | 66.7 | 95.4 | 65.4 |
| CFA | 76.6, 95%CI (64.4,88.8) | 82.4 | 69.2 | 100 | 80.0 | 85.7 | 95.4 | 92.2 |
| PET | 46.8, 95%CI (32.4,61.2) | 70.6 | 26.9 | 75.0 | 43.3 | 80.1 | 90.7 | 71.7 |
| MRI | 55.3, 95%CI (41.0,69.7) | 82.4 | 34.6 | 50.0 | 50.0 | 90.1 | 90.1 | 73.5 |
| CSF | 42.6, 95%CI (28.3,56.8) | 64.7 | 23.1 | 75.0 | 56.7 | 85.7 | 74.4 | 68.6 |
| Age | 40.4, 95%CI (26.2,54.6) | 100 | 7.7 | 0 | 6.7 | 100 | 100 | 50.0 |
| Set of 4 cognitive/functional tests (FAQ, ADAS13, MoCA, MMSE)* | **83.0, 95%CI (72.1,93.8)** | **100** | **69.2** | **100** | **76.7** | **100** | **97.7** | **94.9** |

### 3.2 Development of computer-based decision support tool

Given the high predictive power of CFA and their common use in clinical practice, we developed a prototype of the CDSS for assessing the severity of AD of an individual (based solely on CFA) to aid clinicians to diagnose AD (Fig. 6). The feasibility of our CDSS was demonstrated by using the baseline data from ADNI to benchmark the ability of the AD severity score to model disease prediction. The system implements an automated machine learning approach for data pre-processing, modelling, and validation (as described in Section 2.1) and uses scores of selected cognitive measures as data entries. The disease outcome prediction is generated using the KRR model as it regards the course of disease as a continuous progression and therefore, allows for discriminating between different ‘stages’ of the same AD category (e.g., a light-green colour in Fig. 6 indicates less probable QCI whereas a light-orange colour - more probable QCI). Furthermore, the KRR model achieved the best predictive performance of all regression techniques considered.

**Fig. 6.**
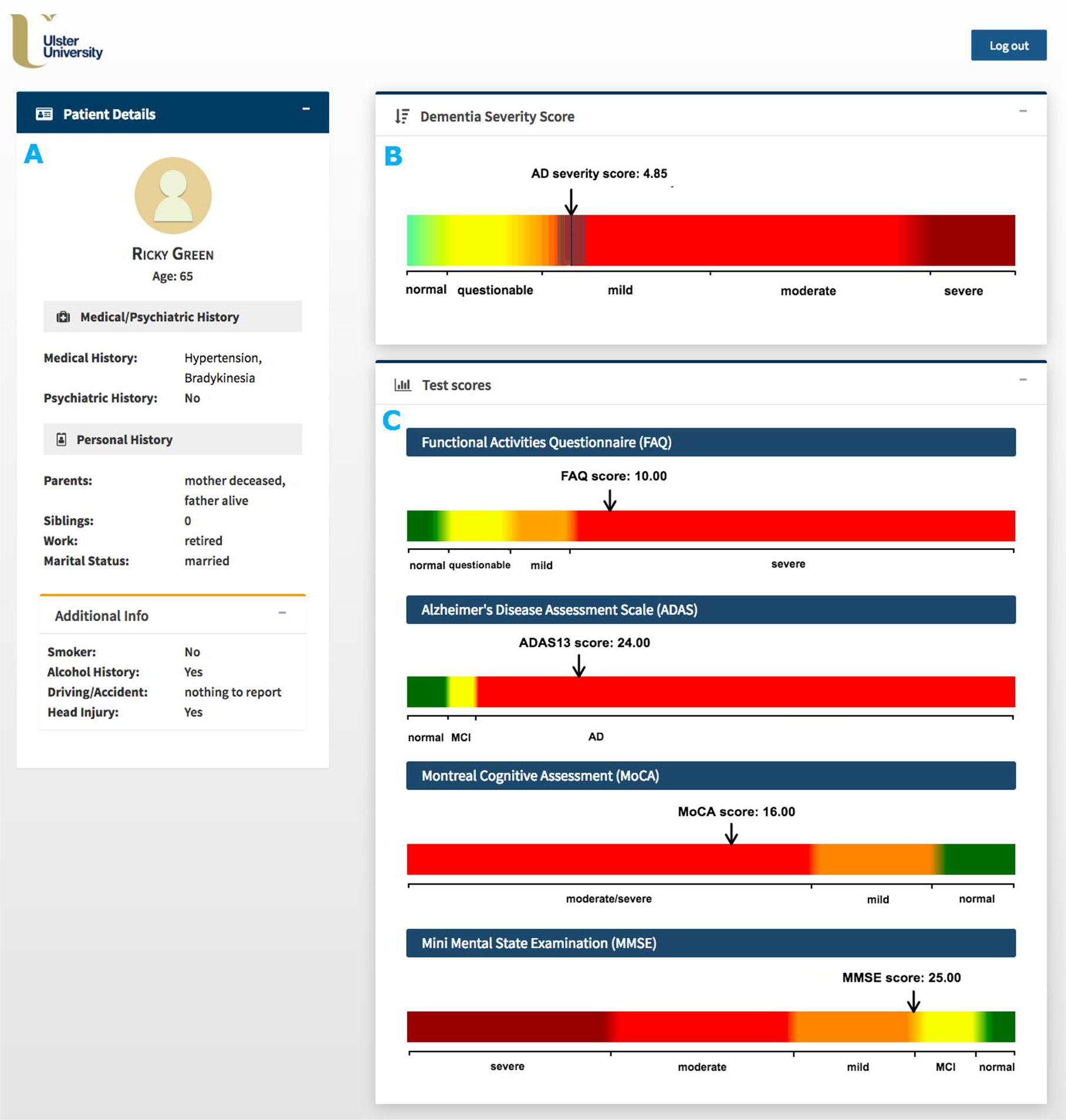
Graphical user interface of the computer-based clinical decision support system for predicting severity of dementia of an individual patient. A) Patient information panel; B) AD severity measurement scale with AD severity score (black line) and its confidence interval (gray range); C) Measurement scales for the selected cognitive/functional assessments (FAQ, ADAS13, MoCA, MMSE). To allow quick interpretation, the AD severity measurement scale is divided into 5 classes based on the CDRSB score, i.e., ‘Normal’ (CDRSB = 0), ‘QCI’ (0.5 ≤ CDRSB ≤ 4.0), ‘AD Mild’ (4.0 ≤ CDRSB ≤ 9), ‘AD Moderate’ (9.5 ≤ CDRSB ≤ 15.5)., and ‘AD severe’ (16 ≤ CDRSB ≤ 18).”

The input panel of our CDSS is designed for a set of 4 CFA inputs, namely, the total scores for FAQ, ADAS13, MoCA, and MMSE. These 4 efficient AD markers achieved the highest performance for the SVM model (MCA of 83%, 95%CI = (72.1%, 93.8%)) while for the KRR model, their performance (*R^2^* = 0.832, 95%CI = (0.754,0.910)) was only slightly lower than best performance reported for the combined CFA and (more labour-intensive and costly) MRI data, i.e., *R^2^* = 0.874, 95%CI = (0.827,0.922) (Table 1 & 2). Although all four tests are commonly used to provide a measure of cognitive impairment in clinical, research, and community settings, they have never been used in combination for evaluating AD severity (Nasreddine et al., 2006, Skinner et al., 2012, Teng et al., 2010, Trzepacz et al., 2015). The MMSE is currently the most widely used screening assessment for general cognitive evaluation and staging of Alzheimer’s disease (Nasreddine et al., 2006, Vertesi et al., 2001). It assesses various cognitive areas including attention, memory, language, orientation, and visuospatial abilities (Vertesi et al., 2001). The MMSE has been frequently applied not only to scale the severity of cognitive impairment at a given point in time but also to document the overall progression of cognitive decline over time (de Souza, Sarazin, Goetz, & Dubois, 2009). When compared to the MMSE, the MoCA consists of more memory, structured language, and executive function items and demonstrates high discriminant potential for MCI patients that performed within the normal range of the MMSE (Nasreddine et al., 2006, Trzepacz et al., 2015, Whitney, Mossbarger, Herman, & Ibarra, 2012). In addition, the MoCA has been shown to exhibit superior sensitivity for amnestic MCI detection compared to the MMSE (Freitas, Simões, Alves, & Santana, 2013). The ADAS13 is mainly applied to evaluate the severity of cognitive and non-cognitive disfunctions from mild to severe AD (Skinner et al., 2012). However, it has also been used as an outcome measure for trials of interventions in people with MCI and appeared to be able to discriminate between patients with MCI and mild AD (Kueper, Speechley, & Montero-Odasso, 2018). In contrast to MMSE, MoCA, and ADAS13, the FAQ is not used in everyday clinical routine (Ritter et al., 2015). However, its relevance for determining impairment in everyday functioning and ensuring accurate early diagnosis of AD has been well-documented (Devanand et al., 2008, Ding et al., 2018, Ritter et al., 2015). For instance, studies found the use of FAQ can significantly contribute to discerning MCI versus AD cases with MoCA scores overlapping in the MCI range (Trzepacz et al., 2015). Furthermore, the FAQ has been shown to be highly sensitive in detecting differences in cognitive functioning between healthy and MCI patients, mainly via the assessment of the ability of assembling documents and remembering appointments (Jekel et al., 2015).

Given the scores of 4 CFA described above, our system is able to provide an evidence-based AD score reflecting the severity of AD in the case of an individual subject. The score is generated by comparing selected CFA scores of an undiagnosed patient against a large database of existing patient records (Figs. 2 & 6). A single patient data with the predicted AD severity score is later added to the clinical data warehouse, updating the database, and initiating the retraining and validation procedure of the predictive model. To highlight the uncertainty inherent in the disease prediction, the system also provides a confidence interval for the predicted AD severity score based on the output from the individual sample validation procedure. Since our approach does not currently use input from clinicians for subsequent learning but uses its own predictions for reinforcing the existing model, further work is required to incorporate a self-training scheme that chooses only high-confidence predictions in the iterative process of model training.

The CDSS patient profile includes only content that is relevant in the context of AD diagnosis, in a concise format to allow quick and unambiguous interpretation. It consists of: 1) the patient information section with patient’s medical, psychiatric, and personal history details (Fig. 6A); 2) the AD severity measurement scale along with the predicted AD score and its confidence interval (Fig. 6B); and 3) CFA test scores together with their corresponding cut-off values for disease classes (Fig. 6C). The AD severity measurement scale is divided into 5 classes based on the CDRSB score i.e. ‘Normal’ (CDRSB = 0), ‘QCI’ (0.5 ≤ CDRSB ≤ 4.0), ‘AD Mild’ (4.0 ≤ CDRSB ≤ 9), ‘AD Moderate’ (9.5 ≤ CDRSB ≤ 15.5)., and ‘AD severe’ (16 ≤ CDRSB ≤ 18). Simple and user-friendly layout of the patient profile allows clinicians to easily assess how different CFA contribute to the predicted AD severity score (Bucholc et al. 2018).

## 4. Discussion

In this study, we have developed a computational framework for identifying key measures in predicting the severity of AD using baseline data from ADNI, which leads to the development of an efficient and practical CDSS prototype for evaluating the severity of AD of an individual on a continuous spectrum. It is efficient in that only a small subset of the data attributes with the highest predictive accuracy of AD severity level is chosen, and they consist of readily available CFA scores. This is practical in the sense that clinical decisions of AD relies relatively heavily on CFA scores. Furthermore, the system uses an automated machine learning approach for data pre-processing, modelling, and validation, making the clinical decision process more objective and accurate.

We showed that model predictions incorporating CFA were more accurate than those based solely on biomarker modalities (single or combinations). The KRR model performed best for the combined CFA and MRI data, i.e., *R^2^* = 0.874, 95%CI = (0.827, 0.922) (Table 1). However the KRR model incorporating only CFA scores (FAQ, ADAS13, MoCA, MMSE) achieved comparable performance, i.e., *R^2^*= 0.832, 95%CI = (0.754, 0.910). The SVR achieved the highest performance for the combination of CFA and MRI, i.e., *R^2^* = 0.790, 95%CI (0.715, 0.866) while kNN_reg_ performed best for CFA, i.e., *R^2^*= 0.750, 95%CI (0.653,0.847) (Table A.2). Given the SVM model, the optimal performance was reported for CFA data, i.e., MCA of 83.0%, 95%CI = (72.1%, 93.8%) for a subset of 4 CFA (FAQ, ADAS13, MoCA, MMSE) (Table 2). Again, the highest accuracy of the RF model was reported for all CFA with MCA of 80.0%, 95%CI (66.7%, 90.9%) while kNN_class_ performed best for the combinations of CFA, MRI and CSF, i.e., MCA of 89.7%, 95%CI (76.9%, 96.5%) (Table A.3, A.4). These results lend support to existing clinical practices that depend relatively heavily on CFAs (Grober, Wakefield, Ehrlich, Mabie & Lipton, 2017). Future analysis of individual tasks making up each of the considered CFAs can lead to building a single optimised CFA.

High predictive power of CFA has been demonstrated in previous studies (Chapman et al., 2011, Cui et al., 2011, Korolev et al., 2016). Cui et al. (2011) showed that single-modality predictive models based on CFA, namely FAQ, LM Delayed Recall, LM Immediate Recall, AVLT Delayed Recall and AVLT trials 1–5 (accuracy of 65%) outperformed those based on volumetric based CSF (accuracy of 60%) and MRI (accuracy of 62%) biomarkers in the task of early identification of MCI patients at risk of progressing to AD. In addition, incorporating multiple data modalities into the model, i.e., CFA, MRI, and CSF data, only slightly improved model performance (accuracy of 67%). Similar observations have been reported by (Chapman et al., 2011, Ewers et al., 2012). Cognitive measures (either alone or combined with other predictors) were also highly predictive in discriminating between stages of cognitive decline (Ewers et al., 2012, Nestor, Scheltens, & Hodges, 2004). In Ewers et al. (2012), the best statistical differentiation between AD and healthy subjects was reached for a combination of neuropsychological tests (RAVLT Immediate and RAVLT Delayed Recall) and CSF t-tau/Aβ_1-4_ratio. However, a single-modality model incorporating cognitive measures showed a predictive accuracy comparable to that of the multi-predictor model. Few other studies claimed relatively good predictive performance of models constructed using tests for memory impairment, abstract reasoning, and verbal fluency (Jacobs et al., 1995, Small, Herlitz, Fratiglioni, Almkvist, & Bäckman, 1997). Note that an increasing number of studies is based on the multimodal approach for either differentiating between stages of disease severity or identifying potential descriptors for the decline of cognition from MCI to AD (Bauer, Cabral, & Killiany, 2018, Ritter et al., 2015). Therefore, it is difficult to assess the individual contributions of modalities, such as CFA, to the accuracy of predictive models. Furthermore, differences in study designs reflected in different data types used, characteristics of patient populations, subject inclusion/exclusion criteria, diagnostic criteria for AD, classification frameworks and evaluation metrics make it challenging to compare results across studies. However, the discriminatory value of cognitive measures in the AD severity assessment or MCI-to-AD conversion has been repeatedly demonstrated.

Numerous predictive approaches have been developed for diagnosis of AD, most of them derived using Cox Regression (Barnes et al., 2014, Derby et al., 2013, Ewers et al., 2012, Okereke et al., 2012, Seshadri et al., 2010), and Logistic Regression (Barnes et al., 2010, Bauer et al., 2018, Chary et al. 2013, Wolfsgruber et al., 2014). In the past decade, there has also been growing interest toward the application of SVM (Casanova et al., 2015, Cui et al., 2011, Klöppel et al., 2008, Ritter et al., 2015, Weygandt et al., 2011), RF (Gray et al., 2013, Sarica et al., 2017) as well as deep neural network models for AD diagnostics (Ortiz, Munilla, Gorriz, & Ramirez, 2016, Shen, Wu, & Suk, 2017). The SVM-based models have been developed for both differential diagnosis and assessment of AD severity using neuroimaging, genome-based, and blood-based biomarkers (Klöppel et al., 2008, Laske et al., 2011, Smith-Vikos & Slack, 2013, Weygandt et al., 2011). RF demonstrated advantages over other ML methods regarding the ability to handle highly non-linearly correlated data (Caruana & Niculescu-Mizil, 2006). While most of deep learning models show great performance in diagnostic classification, their interpretation remains an emerging field of research (Che, Purushotham, Khemani, & Liu, 2016). Other machine learning approaches for assisted diagnosis of cognitive impairment and dementia include linear regression (Agosta et al., 2012, Bauer et al., 2018, Koch et al., 2012), penalized regression (Wang, Liu, & Shen, 2018), Bayesian networks (Ding et al., 2018), hidden Markov models (Wang et al., 2014), and probabilistic multiple kernel learning (MKL) classifiers (Korolev et al., 2016, Youssofzadeh et al., 2017). Despite the common use of machine learning techniques for the disease diagnostics, controversy still exists regarding the effects of different combinations of explanatory variables, hyper-parameter tuning, sample size and class balance on the performance of predictive models (Du, Fu, & Calhoun, 2018, Finch & Schneider, 2007, Michie, Spiegelhalter, & Taylor, 1994). Different applications using different data sets (simulated or real) have failed to generate a model that performed best in all applications (Michie wt al., 1994, Wolpert & MacReady (1997). The results of empirical comparisons often showed opposite results, for example one study claiming that decision trees are superior to neural nets, and another making the opposite claim (Michie et al., 1994). In fact, Wolpert & MacReady (1997) demonstrated the danger of comparing performance of algorithms on a small sample of problems and showed the best learning algorithm is always context dependant.

The integration of efficient, less invasive, and cost-effective clinical markers into CDSS for AD diagnosis of individuals can support prevention-related decision-making in clinical settings. So far, educational interventions aimed at improving GPs’ knowledge and skills in recognition and management of dementia made no significant impact on the number of dementia patients’ care reviews or newly diagnosed cases (Dodd et al., 2015). Despite this, the deployment of CDSSs for a routine use in AD diagnostics, especially those incorporating machine learning methodologies, is still very rare. Furthermore, CDSSs currently used in dementia decision-making require information from expensive and labour-intensive biomarkers (e.g., PredictAD) (Antila et al., 2013) or make use of predictive methodologies based on binary classifications (e.g., CADi2 or CANTAB) (Fray, Robbins & Sahakian, 1996, Onoda & Yamaguchi, 2014). Such approaches are designed to differentiate between two disease categories, e.g., healthy patients and individuals with cognitive impairment. Our computational approach defines the disease in more realistic manner as a continuous progress rather than a sequence of discrete stages. Importantly, it also provides clinician with an estimate of prediction reliability by adopting a validation procedure appropriate for an individual participant data.

Our study has several limitations worth noting. First, our CDSS prototype does not yet include a mechanism for handling missing data. Work is currently in progress to develop an automated approach for missing data imputation that will be later incorporated into the system. Second, the current version of our CDSS provides clinicians with the predicted AD severity score of an individual with the corresponding confidence interval and CFA test scores together with cut-off values for disease classes; however, it does not provide any measures of predictive accuracy of the incorporated model or information regarding the relative importance of individual predictors in the model. We plan to address these issues in future work by making the model evaluation metrics available to clinicians. We also intend to provide the relative importance of individual features incorporated into the model based on the magnitude of standardized regression coefficients. The format of visual representations of performance metrics will be developed in consultation with clinical end-users. Third, the AD measurement scale in our CDSS covers all 5 disease classes i.e. ‘normal’, ‘QCI’, ‘mild’, ‘moderate’, and ‘severe’. However, due to data unavailability, patients with the ‘severe’ type of AD have not been included into our model training set and therefore, such cases could not be learned from the data. The inclusion of the ‘severe’ disease class in the CDSS means the suitability of our KRR model for making predictions outside the range of data used to estimate the model must be further evaluated. The necessary follow-up step would be a testing phase, to establish the degree to which prediction for ‘severe’ cases is contextually valid and hence, clinically useful. This could be done when additional data for patients with the ‘severe’ AD type is obtained. It is also worth noting that the current computational approach implemented into our CDSS is based on the iterative method for semi-supervised learning that uses its own predictions to assign AD severity labels to new (unlabelled) patient data. Accordingly, our CDSS does not use input from clinicians for subsequent learning of the predictive model but uses its own predictions to reinforce the current model. We are aware this might be perceived as the system limitation since the process of training the model on its own predictions may result in model overfitting. Hence, for future work, we plan to enhance our computational framework by incorporating a self-training algorithm for selecting only high-confidence predictions to a training set for the next iteration. Most importantly, we will develop interpretability of our models, either through development of algorithms to “peer” through the black box (Giudotti et al., 2018) or complementing with more interpretable models such as decision trees (Sokol & Flach, 2018). This will facilitate an easy explanation of system’s content and allow for adjustment/correction of the AD severity class based on feedback from clinicians. A dynamic, easily interpretable predictive model interacting with decision makers to re-estimate predictions according to new clinical information could increase the clinical value of our CDSS. Finally, we acknowledge that the proposed CDSS requires further real-time testing and validation in a clinical setting to enhance system’s reliability and stability.

## 5. Conclusion

Our CDSS offers a platform to standardize diagnostics in AD and has the potential to address variations in the quality of GP services associated with the lack of experience or skills in dementia recognition. By taking full advantage of ML techniques, our system can develop, update, and visualize AD risk profiles of individual patients by utilizing only non-invasive and cost-effective AD markers. Although our CDSS has not been designed to provide a diagnosis, it can streamline a clinical workflow and assist with clinical decision-making. As our predictive, ML-based framework becomes more established and its performance better characterized and tested, it could be further upgraded to automate the care pathway for dementia. This process will require the active involvement of the medical community to ensure that developed algorithms are intelligently integrated into existing medical practice and are rigorously validated for clinical efficacy.

## Acknowledgements

Data collection and sharing for this project was funded by the Alzheimer’s Disease Neuroimaging Initiative (ADNI) (National Institutes of Health Grant U01 AG024904) and DOD ADNI (Department of Defense award number W81XWH-12-2-0012). ADNI is funded by the National Institute on Aging, the National Institute of Biomedical Imaging and Bioengineering, and through generous contributions from the following: AbbVie, Alzheimer’s Association; Alzheimer’s Drug Discovery Foundation; Araclon Biotech; BioClinica, Inc.; Biogen; Bristol-Myers Squibb Company; CereSpir, Inc.; Cogstate; Eisai Inc.; Elan Pharmaceuticals, Inc.; Eli Lilly and Company; EuroImmun; F. Hoffmann-La Roche Ltd and its affiliated company Genentech, Inc.; Fujirebio; GE Healthcare; IXICO Ltd.; Janssen Alzheimer Immunotherapy Research & Development, LLC.; Johnson & Johnson Pharmaceutical Research & Development LLC.; Lumosity; Lundbeck; Merck & Co., Inc.; Meso Scale Diagnostics, LLC.; NeuroRx Research; Neurotrack Technologies; Novartis Pharmaceuticals Corporation; Pfizer Inc.; Piramal Imaging; Servier; Takeda Pharmaceutical Company; and Transition Therapeutics. The Canadian Institutes of Health Research is providing funds to support ADNI clinical sites in Canada. Private sector contributions are facilitated by the Foundation for the National Institutes of Health (www.fnih.org). The grantee organization is the Northern California Institute for Research and Education, and the study is coordinated by the Alzheimer’s Therapeutic Research Institute at the University of Southern California. ADNI data are disseminated by the Laboratory for Neuro Imaging at the University of Southern California.

## Funding

This project was previously supported by Innovate UK (102161) (MB, XD, HYW, DHG, HW, GP, LPM, AJB, KWL) and then supported by the EU’s INTERREG VA Programme, managed by the Special EU Programmes Body (SEUPB) (MB, LPM, AJB, PLM, ST, DPF, KWL), the Northern Ireland Functional Brain Mapping Facility (1303/101154803) funded by Invest NI and Ulster University (GP, LPM, AJB, KWL), Ulster University Research Challenge Fund (KWL, XD, PLM, ST), Global Challenges Research Fund (XD, KWL, PLM, ST), and the COST Action Open Multiscale Systems Medicine (OpenMultiMed) supported by COST (European Cooperation in Science and Technology) (KWL). The views and opinions expressed in this paper do not necessarily reflect those of the European Commission or the Special EU Programmes Body (SEUPB).

## Declarations of interest

None

## References

1. Abikoff, H., Alvir, J., Hong, G., Sukoff, R., Orazio, J., Solomon, S. & Saravay, S. (1987). Logical memory subtest of the Wechsler Memory Scale: age and education norms and alternate-form reliability of two scoring systems. Journal of clinical and experimental neuropsychology, 9(4), 435–448.

2. Agosta, F., Pievani, M., Geroldi, C., Copetti, M., Frisoni, G. B., & Filippi, M. (2012). Resting state fMRI in Alzheimer’s disease: beyond the default mode network. Neurobiology of aging, 33(8), 1564–1578.

3. Allen, M. P. (1997). The coefficient of determination in multiple regression. Understanding Regression Analysis, 91–95.

4. Antila, K., Lotjonen, J., Thurfjell, L., Laine, J., Massimini, M., Rueckert, D., Zubarev, R., Oresic, M., van Gils, M., Mattila, J., Hviid Simonsen, A., Waldemar, G. and Soininen, H. (2013). The PredictAD project: development of novel biomarkers and analysis software for early diagnosis of the Alzheimer’s disease. Interface Focus, 3(2), 20120072–20120072.

5. Awad, M., & Khanna, R. (2015). Support vector regression. In Efficient Learning Machines (pp. 67–80). Apress, Berkeley, CA.

6. Barnes, D. E., Beiser, A. S., Lee, A., Langa, K. M., Koyama, A., Preis, S. R., Neuhaus, J., McCammon, R.J., Yaffe, K., Seshadri, S. & Haan, M. N. (2014). Development and validation of a brief dementia screening indicator for primary care. Alzheimer’s & Dementia, 10(6), 656–665.

7. Barnes, D. E., Covinsky, K. E., Whitmer, R. A., Kuller, L. H., Lopez, O. L., & Yaffe, K. (2010). Dementia risk indices: A framework for identifying individuals with a high dementia risk. Alzheimer’s & Dementia, 6(2), 138.

8. Basak, D., Pal, S., & Patranabis, D. C. (2007). Support vector regression. Neural Information Processing-Letters and Reviews, 11(10), 203–224.

9. Bauer, C. M., Cabral, H. J., & Killiany, R. J. (2018). Multimodal Discrimination between Normal Aging, Mild Cognitive Impairment and Alzheimer’s Disease and Prediction of Cognitive Decline. Diagnostics, 8(1), 14.

10. Bengio, Y., Delalleau, O., Roux, N. L., Paiement, J. F., Vincent, P., & Ouimet, M. (2003). Feature Extraction: Foundations and Applications, chapter Spectral Dimensionality Reduction. Springer.

11. Benoít, F., Van Heeswijk, M., Miche, Y., Verleysen, M., & Lendasse, A. (2013). Feature selection for nonlinear models with extreme learning machines. Neurocomputing, 102, 111–124.

12. Brodaty, H., Woolf, C., Andersen, S., Barzilai, N., Brayne, C., Cheung, K., Corrada, M., Crawford, J., Daly, C., Gondo, Y., Hagberg, B., Hirose, N., Holstege, H., Kawas, C., Kaye, J., Kochan, N., Lau, B., Lucca, U., Marcon, G., Martin, P., Poon, L., Richmond, R., Robine, J., Skoog, I., Slavin, M., Szewieczek, J., Tettamanti, M., Viña, J., Perls, T. and Sachdev, P. (2016). ICC-dementia (International Centenarian Consortium - dementia): an international consortium to determine the prevalence and incidence of dementia in centenarians across diverse ethnoracial and sociocultural groups. BMC Neurology, 16(1).

13. Brown, S. A. (2016). Patient similarity: emerging concepts in systems and precision medicine. Frontiers in physiology, 7, 561.

14. Bucholc, M.*, Ding, X.*, Wang, H.Y., Glass, D., Wang, H., Prasad, G., Maguire, L.P., Bjourson, A.J., McClean, P.L., Todd, S., Finn, D.P. & Wong-Lin, K. (2018, March). Development of a computer-based clinical decision support tool for identifying individuals with different levels of cognitive impairment. Poster session presentation at the meeting of Alzheimer’s Research UK, London. *\*Joint first authors*.

15. Bucholc, M., Ding, X., Wang, H., Glass, D.H., Wang, H., Bjourson, A.J., Dowey, LR., O’Kane, M., Maguire, L., Prasad, G. & Wong-Lin, K. (2017, September). Data analytics and computerised application for predicting Alzheimer’s disease severity and related outlier test scores, Poster session presentation at the meeting of Translational Medicine (TMED) 8 Conference, Derry-Londonderry.

16. Caruana, R., & Niculescu-Mizil, A. (2006). An empirical comparison of supervised learning algorithms. In Proceedings of the 23rd international conference on Machine learning (pp. 161–168). ACM.

17. Casanova, R., Hsu, F. C., Espeland, M. A., & Alzheimer’s Disease Neuroimaging Initiative. (2012). Classification of structural MRI images in Alzheimer’s disease from the perspective of ill-posed problems. PloS one, 7(10), e44877.

18. Castaneda, C., Nalley, K., Mannion, C., Bhattacharyya, P., Blake, P., Pecora, A., Goy, A. & Suh, K. S. (2015). Clinical decision support systems for improving diagnostic accuracy and achieving precision medicine. Journal of clinical bioinformatics, 5(1), 4.

19. Cedarbaum, J. M., Jaros, M., Hernandez, C., Coley, N., Andrieu, S., Grundman, M., Vellas, B. & Alzheimer’s Disease Neuroimaging Initiative. (2013). Rationale for use of the Clinical Dementia Rating Sum of Boxes as a primary outcome measure for Alzheimer’s disease clinical trials. Alzheimer’s & Dementia, 9(1), S45–S55.

20. Chapman, R. M., Mapstone, M., McCrary, J. W., Gardner, M. N., Porsteinsson, A., Sandoval, T. C., Guillily, M.D., DeGrush, E. & Reilly, L. A. (2011). Predicting conversion from mild cognitive impairment to Alzheimer’s disease using neuropsychological tests and multivariate methods. Journal of Clinical and Experimental Neuropsychology, 33(2), 187–199.

21. Chary, E., Amieva, H., Pérès, K., Orgogozo, J. M., Dartigues, J. F., & Jacqmin-Gadda, H. (2013). Short-versus long-term prediction of dementia among subjects with low and high educational levels. Alzheimer’s & Dementia, 9(5), 562–571.

22. Chawla, N. V., Bowyer, K. W., Hall, L. O., & Kegelmeyer, W. P. (2002). SMOTE: synthetic minority over-sampling technique. Journal of artificial intelligence research, 16, 321–357.

23. Che, Z., Purushotham, S., Khemani, R., & Liu, Y. (2016). Interpretable deep models for icu outcome prediction. In AMIA Annual Symposium Proceedings (Vol. 2016, p. 371). American Medical Informatics Association.

24. Chen, K., Langbaum, J. B., Fleisher, A. S., Ayutyanont, N., Reschke, C., Lee, W., Liu, X., Bandy, D., Alexander, G.E., Thompson, P.M. & Foster, N. L. (2010). Twelve-month metabolic declines in probable Alzheimer’s disease and amnestic mild cognitive impairment assessed using an empirically pre-defined statistical region-of-interest: findings from the Alzheimer’s Disease Neuroimaging Initiative. Neuroimage, 51(2), 654–664.

25. Cortes, C., & Vapnik, V. (1995). Support-vector networks. Machine learning, 20(3), 273–297.

26. Cortizo, J. C., & Giraldez, I. (2006). Multi criteria wrapper improvements to naive bayes learning. In International Conference on Intelligent Data Engineering and Automated Learning (pp. 419–427). Springer, Berlin.

27. Cui, Y., Liu, B., Luo, S., Zhen, X., Fan, M., Liu, T., Zhu, W., Park, M., Jiang, T., Jin, J.S. & Alzheimer’s Disease Neuroimaging Initiative. (2011). Identification of conversion from mild cognitive impairment to Alzheimer’s disease using multivariate predictors. PloS one, 6(7), e21896.

28. Dagliati, A., Tibollo, V., Sacchi, L., Malovini, A., Limongelli, I., Gabetta, M., Napolitano, C., Mazzanti, A., De Cata, P., Chiovato, L. & Priori, S. (2018). Big Data as a driver for Clinical Decision Support Systems: a Learning Health Systems perspective. Frontiers in Digital Humanities, 5, 8.

29. de Souza, L. C., Sarazin, M., Goetz, C., & Dubois, B. (2009). Clinical investigations in primary care. Dementia in Clinical Practice, 24, 1–11.

30. Devanand, D. P., Liu, X., Tabert, M. H., Pradhaban, G., Cuasay, K., Bell, K., de Leon, M.J., Doty, R.L., Stern, Y. & Pelton, G. H. (2008). Combining early markers strongly predicts conversion from mild cognitive impairment to Alzheimer’s disease. Biological psychiatry, 64(10), 871–879.

31. Derby, C. A., Burns, L. C., Wang, C., Katz, M. J., Zimmerman, M. E., L’italien, G., Guo, Z., Berman, R.M. & Lipton, R. B. (2013). Screening for predementia AD time-dependent operating characteristics of episodic memory tests. Neurology, 80(14), 1307–1314.

32. Ding, X., Bucholc, M., Wang, H., Glass, D. H., Wang, H., Clarke, D. H., Bjourson, A.J., Le Roy, C.D., O’Kane, M., Prasad, G. & Maguire, L. & Wong-Lin, K. (2018). A hybrid computational approach for efficient Alzheimer’s disease classification based on heterogeneous data. Scientific reports, 8(1), 9774.

33. Dodd, E., Cheston, R., & Ivanecka, A. (2015). The assessment of dementia in primary care. Journal of psychiatric and mental health nursing, 22(9), 731–737.

34. Du, Y., Fu, Z., & Calhoun, V. D. (2018). Classification and prediction of brain disorders using functional connectivity: Promising but challenging. Frontiers in neuroscience, 12.

35. Duchesne, S., Caroli, A., Geroldi, C., Collins, D. L., & Frisoni, G. B. (2009). Relating one-year cognitive change in mild cognitive impairment to baseline MRI features. Neuroimage, 47(4), 1363–1370.

36. Duchesne, S., Caroli, A., Geroldi, C., Frisoni, G. B., & Collins, D. L. (2005, October). Predicting clinical variable from MRI features: application to MMSE in MCI. In International Conference on Medical Image Computing and Computer-Assisted Intervention (pp. 392–399). Springer, Berlin, Heidelberg.

37. Dyrba, M., Grothe, M., Kirste, T., & Teipel, S. J. (2015). Multimodal analysis of functional and structural disconnection in A lzheimer’s disease using multiple kernel SVM. Human brain mapping, 36(6), 2118–2131.

38. Elisseeff, A., & Pontil, M. (2003). Leave-one-out error and stability of learning algorithms with applications. NATO science series sub series iii computer and systems sciences, 190, 111–130.

39. Ewers, M., Walsh, C., Trojanowski, J. Q., Shaw, L. M., Petersen, R. C., Jack Jr, C. R., Feldman, H.H., Bokde, A.L., Alexander, G.E., Scheltens, P. & Vellas, B. (2012). Prediction of conversion from mild cognitive impairment to Alzheimer’s disease dementia based upon biomarkers and neuropsychological test performance. Neurobiology of aging, 33(7), 1203–1214.

40. Farran, B., Channanath, A. M., Behbehani, K., & Thanaraj, T. A. (2013). Predictive models to assess risk of type 2 diabetes, hypertension and comorbidity: machine-learning algorithms and validation using national health data from Kuwait—a cohort study. BMJ open, 3(5), e002457.

41. Finch, H., & Schneider, M. K. (2007). Classification accuracy of neural networks vs. discriminant analysis, logistic regression, and classification and regression trees. Methodology, 3(2), 47–57.

42. Folstein, M. F., Robins, L. N., & Helzer, J. E. (1983). The mini-mental state examination. Archives of general psychiatry, 40(7), 812–812.

43. Forlenza, O. V., Radanovic, M., Talib, L. L., Aprahamian, I., Diniz, B. S., Zetterberg, H., & Gattaz, W. F. (2015). Cerebrospinal fluid biomarkers in Alzheimer’s disease: Diagnostic accuracy and prediction of dementia. Alzheimer’s & Dementia: Diagnosis, Assessment & Disease Monitoring, 1(4), 455–463.

44. Frame, A., LaMantia, M., Bynagari, B. B. R., Dexter, P., & Boustani, M. (2013). Development and implementation of an electronic decision support to manage the health of a high-risk population: the enhanced Electronic Medical Record Aging Brain Care Software (eMR-ABC). EGEMS, 1(1).

45. Fray, J. P., Robbins, W. T., & Sahakian, J. B. (1996). Neuorpsychiatyric applications of CANTAB. International journal of geriatric psychiatry, 11(4), 329–336.

46. Freitas, S., Simões, M. R., Alves, L., & Santana, I. (2013). Montreal cognitive assessment: validation study for mild cognitive impairment and Alzheimer disease. Alzheimer Disease & Associated Disorders, 27(1), 37–43.

47. Gálvez, J. A., Ahumada, L., Simpao, A. F., Lin, E. E., Bonafide, C. P., Choudhry, D., England, W.R., Jawad, A.F., Friedman, D., Sesok-Pizzini, D.A. & Rehman, M. A. (2013). Visual analytical tool for evaluation of 10-year perioperative transfusion practice at a children’s hospital. Journal of the American Medical Informatics Association, 21(3), 529–534.

48. Granitto, P. M., Furlanello, C., Biasioli, F., & Gasperi, F. (2006). Recursive feature elimination with random forest for PTR-MS analysis of agroindustrial products. Chemometrics and Intelligent Laboratory Systems, 83(2), 83–90.

49. Gregorutti, B., Michel, B., & Saint-Pierre, P. (2017). Correlation and variable importance in random forests. Statistics and Computing, 27(3), 659–678.

50. Grimmer, T., Wutz, C., Alexopoulos, P., Drzezga, A., Förster, S., Förstl, H., Goldhardt, O., Ortner, M., Sorg, C. & Kurz, A. (2016). Visual versus fully automated analyses of 18F-FDG and amyloid PET for prediction of dementia due to Alzheimer disease in mild cognitive impairment. Journal of Nuclear Medicine, 57(2), 204–207.

51. Grober, E., Wakefield, D., Ehrlich, A. R., Mabie, P., & Lipton, R. B. (2017). Identifying memory impairment and early dementia in primary care. Alzheimer’s & Dementia: Diagnosis, Assessment & Disease Monitoring, 6, 188–195.

52. Guidotti, R., Monreale, A., Ruggieri, S., Turini, F., Giannotti, F., & Pedreschi, D. (2018). A survey of methods for explaining black box models. ACM Computing Surveys (CSUR), 51(5), 93.

53. Hainmueller, J., & Hazlett, C. (2014). Kernel regularized least squares: Reducing misspecification bias with a flexible and interpretable machine learning approach. Political Analysis, 22(2), 143–168.

54. Hand, D. J., & Till, R. J. (2001). A simple generalisation of the area under the ROC curve for multiple class classification problems. Machine learning, 45(2), 171–186.

55. Handels, R.L., Vos, S.J., Kramberger, M.G., Jelic, V., Blennow, K., van Buchem, M., van der Flier, W., Freund-Levi, Y., Hampel, H., Rikkert, M.O. & Oleksik, A. (2017). Predicting progression to dementia in persons with mild cognitive impairment using cerebrospinal fluid markers. Alzheimer’s & Dementia, 13(8), 903–912.

56. Helldén, A., Al-Aieshy, F., Bastholm-Rahmner, P., Bergman, U., Gustafsson, L. L., Höök, H., Sjöviker, S., Söderström, A. & Odar-Cederlöf, I. (2015). Development of a computerised decisions support system for renal risk drugs targeting primary healthcare. BMJ open, 5(7), e006775.

57. Higdon, R., Foster, N. L., Koeppe, R. A., DeCarli, C. S., Jagust, W. J., Clark, C. M., Barbas, N.R., Arnold, S.E., Turner, R.S., Heidebrink, J.L. & Minoshima, S. (2004). A comparison of classification methods for differentiating fronto-temporal dementia from Alzheimer’s disease using FDG-PET imaging. Statistics in medicine, 23(2), 315–326.

58. Hira, Z. M., & Gillies, D. F. (2015). A review of feature selection and feature extraction methods applied on microarray data. Advances in bioinformatics, 2015.

59. Jacobs, D. M., Sano, M., Dooneief, G., Marder, K., Bell, K. L., & Stern, Y. (1995). Neuropsychological detection and characterization of preclinical Alzheimer’s disease. Neurology, 45(5), 957–962.

60. Jagust, W. J., Bandy, D., Chen, K., Foster, N. L., Landau, S. M., Mathis, C. A., Price, J.C., Reiman, E.M., Skovronsky, D., Koeppe, R.A. & Alzheimer’s Disease Neuroimaging Initiative. (2010). The Alzheimer’s Disease Neuroimaging Initiative positron emission tomography core. Alzheimer’s & Dementia, 6(3), 221–229.

61. Jekel, K., Damian, M., Wattmo, C., Hausner, L., Bullock, R., Connelly, P. J., Dubois, B., Eriksdotter, M., Ewers, M., Graessel, E. & Kramberger, M. G. (2015). Mild cognitive impairment and deficits in instrumental activities of daily living: a systematic review. Alzheimer’s research & therapy, 7(1), 17.

62. Karas, G., Sluimer, J., Goekoop, R., Van Der Flier, W., Rombouts, S. A. R. B., Vrenken, H., Scheltens, P., Fox, N. & Barkhof, F. (2008). Amnestic mild cognitive impairment: structural MR imaging findings predictive of conversion to Alzheimer disease. American Journal of Neuroradiology, 29(5), 944–949.

63. Klöppel, S., Stonnington, C. M., Chu, C., Draganski, B., Scahill, R. I., Rohrer, J. D., Fox, N.C., Jack Jr, C.R., Ashburner, J. & Frackowiak, R. S. (2008). Automatic classification of MR scans in Alzheimer’s disease. Brain, 131(3), 681–689.

64. Koch, T., Iliffe, S. & EVIDEM-ED project (2010). Rapid appraisal of barriers to the diagnosis and management of patients with dementia in primary care: a systematic review. BMC family practice, 11(1), 52.

65. Koch, W., Teipel, S., Mueller, S., Benninghoff, J., Wagner, M., Bokde, A. L., Hampel, H., Coates, U., Reiser, M. & Meindl, T. (2012). Diagnostic power of default mode network resting state fMRI in the detection of Alzheimer’s disease. Neurobiology of aging, 33(3), 466–478.

66. Korolev, I. O., Symonds, L. L., Bozoki, A. C., & Alzheimer’s Disease Neuroimaging Initiative. (2016). Predicting progression from mild cognitive impairment to Alzheimer’s dementia using clinical, MRI, and plasma biomarkers via probabilistic pattern classification. PloS one, 11(2), e0138866.

67. Kramer, O. (2013). Dimensionality reduction with unsupervised nearest neighbors. Berlin Heidelberg: Springer.

68. Kueper, J. K., Speechley, M., & Montero-Odasso, M. (2018). The Alzheimer’s Disease Assessment Scale–Cognitive Subscale (ADAS-Cog): Modifications and Responsiveness in Pre-Dementia Populations. A Narrative Review. Journal of Alzheimer’s Disease, (Preprint), 1–22.

69. Lama, R. K., Gwak, J., Park, J. S., & Lee, S. W. (2017). Diagnosis of Alzheimer’s disease based on structural MRI images using a regularized extreme learning machine and PCA features. Journal of healthcare engineering, 2017.

70. Landau, S. M., Harvey, D., Madison, C. M., Koeppe, R. A., Reiman, E. M., Foster, N. L., Weiner, M.W., Jagust, W.J. & Alzheimer’s Disease Neuroimaging Initiative. (2011). Associations between cognitive, functional, and FDG-PET measures of decline in AD and MCI. Neurobiology of aging, 32(7), 1207–1218.

71. Landau, S. M., Mintun, M. A., Joshi, A. D., Koeppe, R. A., Petersen, R. C., Aisen, P. S., Weiner, M.W., Jagust, W.J. & Alzheimer’s Disease Neuroimaging Initiative. (2012). Amyloid deposition, hypometabolism, and longitudinal cognitive decline. Annals of neurology, 72(4), 578–586.

72. Lang, L., Clifford, A., Wei, L., Zhang, D., Leung, D., Augustine, G., Danat, I.M., Zhou, W., Copeland, J.R., Anstey, K.J. & Chen, R. (2017). Prevalence and determinants of undetected dementia in the community: a systematic literature review and a meta-analysis. BMJ open, 7(2), e011146.

73. Laske, C., Leyhe, T., Stransky, E., Hoffmann, N., Fallgatter, A. J., & Dietzsch, J. (2011). Identification of a blood-based biomarker panel for classification of Alzheimer’s disease. International Journal of Neuropsychopharmacology, 14(9), 1147–1155.

74. Lebedeva, A. K., Westman, E., Borza, T., Beyer, M. K., Engedal, K., Aarsland, D., Selbaek, G. & Haberg, A. K. (2017). MRI-based classification models in prediction of mild cognitive impairment and dementia in late-life depression. Frontiers in aging neuroscience, 9, 13.

75. Li, Z., Xie, W., & Liu, T. (2018). Efficient feature selection and classification for microarray data. PloS one, 13(8), e0202167.

76. Lindgren, H. (2011). Towards personalized decision support in the dementia domain based on clinical practice guidelines. User Modeling and User-Adapted Interaction, 21(4-5), 377–406.

77. Lindgren, H., Eklund, P., & Eriksson, S. (2002). Clinical decision support system in dementia care. Studies in health technology and informatics, 90, 568–571.

78. Lindquist, A. M., Johansson, P. E., Petersson, G. I., Saveman, B. I., & Nilsson, G. C. (2008). The use of the Personal Digital Assistant (PDA) among personnel and students in health care: a review. Journal of medical Internet research, 10(4).

79. Lisboa, P. J., & Taktak, A. F. (2006). The use of artificial neural networks in decision support in cancer: a systematic review. Neural networks, 19(4), 408–415.

80. Liu, X., Cao, P., Yang, J., & Zhao, D. (2018). Linearized and Kernelized Sparse Multitask Learning for Predicting Cognitive Outcomes in Alzheimer’s Disease. Computational and mathematical methods in medicine, 2018.

81. Liu, H., & Motoda, H. (Eds.). (2007). Computational methods of feature selection. CRC Press.

82. Long, X., Chen, L., Jiang, C., Zhang, L., & Alzheimer’s Disease Neuroimaging Initiative. (2017). Prediction and classification of Alzheimer disease based on quantification of MRI deformation. PloS one, 12(3), e0173372.

83. Magnin, B., Mesrob, L., Kinkingnéhun, S., Pélégrini-Issac, M., Colliot, O., Sarazin, M., Dubois, B., Lehéricy, S. & Benali, H. (2009). Support vector machine-based classification of Alzheimer’s disease from whole-brain anatomical MRI. Neuroradiology, 51(2), 73–83.

84. Maldonado, S., & Weber, R. (2009). A wrapper method for feature selection using support vector machines. Information Sciences, 179(13), 2208–2217.

85. Maldonado, S., Weber, R., & Famili, F. (2014). Feature selection for high-dimensional class-imbalanced data sets using Support Vector Machines. Information Sciences, 286, 228–246.

86. Mandala, P. K., Saharana, S., Khana, S. A., & Jamesa, M. (2015). Apps for dementia screening: a cost-effective and portable solution. Journal of Alzheimer’s Disease, 47(4), 869–872.

87. Maroco, J., Silva, D., Rodrigues, A., Guerreiro, M., Santana, I., & de Mendonça, A. (2011). Data mining methods in the prediction of Dementia: A real-data comparison of the accuracy, sensitivity and specificity of linear discriminant analysis, logistic regression, neural networks, support vector machines, classification trees and random forests. BMC research notes, 4(1), 299.

88. Matheny, M. E., Resnic, F. S., Arora, N., & Ohno-Machado, L. (2007). Effects of SVM parameter optimization on discrimination and calibration for post-procedural PCI mortality. Journal of Biomedical Informatics, 40(6), 688–697.

89. Mattsson, N., Zetterberg, H., Hansson, O., Andreasen, N., Parnetti, L., Jonsson, M., Herukka, S.K., van der Flier, W.M., Blankenstein, M.A., Ewers, M. & Rich, K. (2009). CSF biomarkers and incipient Alzheimer disease in patients with mild cognitive impairment. Jama, 302(4), 385–393.

90. Michalak, K., & Kwaśnicka, H. (2006). Correlation-based feature selection strategy in classification problems. International Journal of Applied Mathematics and Computer Science, 16, 503–511.

91. Mohs, R. C., Knopman, D., Petersen, R. C., Ferris, S. H., Ernesto, C., Grundman, M., Sano, M., Bieliauskas, L., Geldmacher, D., Clark, C. & Thal, L. J. (1997). Development of cognitive instruments for use in clinical trials of antidementia drugs: additions to the Alzheimer’s Disease Assessment Scale that broaden its scope. Alzheimer disease and associated disorders, 11, 13–21.

92. Moja, L., Friz, H. P., Capobussi, M., Kwag, K., Banzi, R., Ruggiero, F., González-Lorenzo, M., Liberati, E.G., Mangia, M., Nyberg, P. & Kunnamo, I. (2015). Implementing an evidence-based computerized decision support system to improve patient care in a general hospital: the CODES study protocol for a randomized controlled trial. Implementation Science, 11(1), 89.

93. Moradi, E., Pepe, A., Gaser, C., Huttunen, H., Tohka, J., & Alzheimer’s Disease Neuroimaging Initiative. (2015). Machine learning framework for early MRI-based Alzheimer’s conversion prediction in MCI subjects. Neuroimage, 104, 398–412.

94. Murphy, K. P. (2014). Machine learning, a probabilistic perspective. MIT Press.

95. Nasreddine, Z. S., Phillips, N. A., Bédirian, V., Charbonneau, S., Whitehead, V., Collin, I., Cummings, J.L. & Chertkow, H. (2005). The Montreal Cognitive Assessment, MoCA: a brief screening tool for mild cognitive impairment. Journal of the American Geriatrics Society, 53(4), 695–699.

96. Nestor, P. J., Scheltens, P., & Hodges, J. R. (2004). Advances in the early detection of Alzheimer’s disease. Nature medicine, 10(7), S34.

97. Okereke, O. I., Pantoja-Galicia, N., Copeland, M., Hyman, B. T., Wanggaard, T., Albert, M. S., Betensky, R.A. & Blacker, D. (2012). The SIST-M: predictive validity of a brief structured clinical dementia rating interview. Alzheimer disease and associated disorders, 26(3), 225.

98. Onoda, K., & Yamaguchi, S. (2014). Revision of the cognitive assessment for dementia, iPad Version (CADi2). PloS one, 9(10), e109931.

99. Ortiz, A., Munilla, J., Gorriz, J. M., & Ramirez, J. (2016). Ensembles of deep learning architectures for the early diagnosis of the Alzheimer’s disease. International journal of neural systems, 26(07), 1650025.

100. Panthong, R., & Srivihok, A. (2015). Wrapper feature subset selection for dimension reduction based on ensemble learning algorithm. Procedia Computer Science, 72, 162–169.

101. Paterson, N. E., & Pond, D. (2009). Early diagnosis of dementia in primary care in Australia: a qualitative study into the barriers and enablers. Alzheimer’s & Dementia: The Journal of the Alzheimer’s Association, 5(4), P185.

102. Perez-Riverol, Y., Kuhn, M., Vizcaíno, J. A., Hitz, M. P., & Audain, E. (2017). Accurate and fast feature selection workflow for high-dimensional omics data. PloS one, 12(12), e0189875.

103. Pfeffer, R. I., Kurosaki, T. T., Harrah Jr, C. H., Chance, J. M., & Filos, S. (1982). Measurement of functional activities in older adults in the community. Journal of gerontology, 37(3), 323–329.

104. Ramírez, J., Górriz, J. M., Salas-Gonzalez, D., Romero, A., López, M., Álvarez, I., & Gómez-Río, M. (2013). Computer-aided diagnosis of Alzheimer’s type dementia combining support vector machines and discriminant set of features. Information Sciences, 237, 59–72. Rey, A. (1964). The clinical examination in psychology. Paris: Presses Universitaires de France.

105. Ritchie, C., Russ, T., Banerjee, S., Barber, B., Boaden, A., Fox, N., Holmes, C., Isaacs, J., Leroi, I., Lovestone, S., Norton, M., O’Brien, J., Pearson, J., Perry, R., Pickett, J., Waldman, A., Wong, W., Rossor, M. and Burns, A. (2017). The Edinburgh Consensus: preparing for the advent of disease-modifying therapies for Alzheimer’s disease. Alzheimer’s Research & Therapy, 9(1), 85.

106. Ritter, K., Schumacher, J., Weygandt, M., Buchert, R., Allefeld, C., Haynes, J. D., & Alzheimer’s Disease Neuroimaging Initiative. (2015). Multimodal prediction of conversion to Alzheimer’s disease based on incomplete biomarkers. Alzheimer’s & Dementia: Diagnosis, Assessment & Disease Monitoring, 1(2), 206–215.

107. Rodin, A. S., Litvinenko, A., Klos, K., Morrison, A. C., Woodage, T., Coresh, J., & Boerwinkle, E. (2009). Use of wrapper algorithms coupled with a random forests classifier for variable selection in large-scale genomic association studies. Journal of computational biology, 16(12), 1705–1718.

108. Samper-González, J., Burgos, N., Bottani, S., Fontanella, S., Lu, P., Marcoux, A., Routier, A., Guillon, J., Bacci, M., Wen, J., Bertrand, A., Bertin, H., Habert, M., Durrleman, S., Evgeniou, T., Colliot, O., Alzheimer’s Disease Neuroimaging Initiative & Australian Imaging Biomarkers and Lifestyle flagship study of ageing (2018). Reproducible evaluation of classification methods in Alzheimer’s disease: framework and application to MRI and PET data. Neuroimage, 183, 504–521.

109. Sanchez-Catasus, C.A., Stormezand, G.N., Jan van Laar, P., De Deyn, P., Alvarez Sanchez, M., & Dierckx, R. (2017). FDG-PET for prediction of AD dementia in mild cognitive impairment. A review of the state of the art with particular emphasis on the comparison with other neuroimaging modalities (MRI and perfusion SPECT). Current Alzheimer Research, 14(2), 127–142.

110. Sarica, A., Cerasa, A., & Quattrone, A. (2017). Random Forest Algorithm for the Classification of Neuroimaging Data in Alzheimer’s Disease: A Systematic Review. Frontiers in Aging Neuroscience, 9, 329.

111. Seshadri, S., Fitzpatrick, A. L., Ikram, M. A., DeStefano, A. L., Gudnason, V., Boada, M., Bis, J.C., Smith, A.V., Carrasquillo, M.M., Lambert, J.C. & Harold, D. (2010). Genome-wide analysis of genetic loci associated with Alzheimer disease. Jama, 303(18), 1832–1840.

112. Shen, D., Wu, G., & Suk, H. I. (2017). Deep learning in medical image analysis. Annual review of biomedical engineering, 19, 221–248.

113. Skinner, J., Carvalho, J. O., Potter, G. G., Thames, A., Zelinski, E., Crane, P. K., Gibbons, L.E. & Alzheimer’s Disease Neuroimaging Initiative. (2012). The Alzheimer’s disease assessment scale-cognitive-plus (ADAS-Cog-Plus): an expansion of the ADAS-Cog to improve responsiveness in MCI. Brain imaging and behavior, 6(4), 489–501.

114. Skyttberg, N., Vicente, J., Chen, R., Blomqvist, H., & Koch, S. (2016). How to improve vital sign data quality for use in clinical decision support systems? A qualitative study in nine Swedish emergency departments. BMC medical informatics and decision making, 16(1), 61.

115. Small, B. J., Herlitz, A., Fratiglioni, L., Almkvist, O., & Bäckman, L. (1997). Cognitive predictors of incident Alzheimer’s disease: a prospective longitudinal study. Neuropsychology, 11(3), 413.

116. Smith-Vikos, T. & Slack, F.J. (2013). MicroRNAs circulate around Alzheimer’s disease. Genome Biol. 14, 125.

117. Sokol, K., & Flach, P. A. (2018). Glass-Box: Explaining AI Decisions With Counterfactual Statements Through Conversation With a Voice-enabled Virtual Assistant. In IJCAI (pp. 5868–5870).

118. Soininen, H., Mattila, J., Koikkalainen, J., van Gils, M., Hviid Simonsen, A., Waldemar, G., Rueckert, D., Thurfjell, L. and Lötjönen, J. (2012). Software Tool for Improved Prediction of Alzheimer’s Disease. Neurodegenerative Diseases, 10(1-4), 149–152.

119. Teipel, S. J., Grothe, M. J., Metzger, C. D., Grimmer, T., Sorg, C., Ewers, M., Franzmeier, N., Meisenzahl, E., Klöppel, S., Borchardt, V. & Walter, M. (2017). Robust detection of impaired resting state functional connectivity networks in alzheimer’s disease using elastic net regularized regression. Frontiers in aging neuroscience, 8, 318.

120. Teng, E., Becker, B. W., Woo, E., Knopman, D. S., Cummings, J. L., & Lu, P. H. (2010). Utility of the Functional Activities Questionnaire for distinguishing mild cognitive impairment from very mild Alzheimer’s disease. Alzheimer disease and associated disorders, 24(4), 348.

121. Tripoliti, E. E., Fotiadis, D. I., Argyropoulou, M., & Manis, G. (2010). A six stage approach for the diagnosis of the Alzheimer’s disease based on fMRI data. Journal of biomedical informatics, 43(2), 307–320.

122. Trzepacz, P. T., Hochstetler, H., Wang, S., Walker, B., Saykin, A. J., Alzheimer’s Disease Neuroimaging Initiative (2015). Relationship between the Montreal Cognitive Assessment and Mini-mental State Examination for assessment of mild cognitive impairment in older adults. BMC geriatrics, 15, 107.

123. Vertesi, A., Lever, J. A., Molloy, D. W., Sanderson, B., Tuttle, I., Pokoradi, L., & Principi, E. (2001). Standardized Mini-Mental State Examination. Use and interpretation. Canadian Family Physician, 47(10), 2018–2023.

124. Vu, K., Snyder, J. C., Li, L., Rupp, M., Chen, B. F., Khelif, T., Muller, K. R. & Burke, K. (2015). Understanding kernel ridge regression: Common behaviors from simple functions to density functionals. International Journal of Quantum Chemistry, 115(16), 1115–1128.

125. Wang, P., Liu, Y., & Shen, D. (2018). Flexible Locally Weighted Penalized Regression with Applications on Prediction of Alzheimer’s Disease Neuroimaging Initiative’s Clinical Scores. IEEE transactions on medical imaging.

126. Wang, Y., Resnick, S. M., Davatzikos, C., Baltimore Longitudinal Study of Aging, & Alzheimer’s Disease Neuroimaging Initiative. (2014). Analysis of spatio-temporal brain imaging patterns by hidden markov models and serial MRI images. Human brain mapping, 35(9), 4777–4794.

127. Weygandt, M., Hackmack, K., Pfüller, C., Bellmann-Strobl, J., Paul, F., Zipp, F., & Haynes, J. D. (2011). MRI pattern recognition in multiple sclerosis normal-appearing brain areas. PLoS One, 6(6), e21138.

128. Whitney, K. A., Mossbarger, B., Herman, S. M., & Ibarra, S. L. (2012). Is the Montreal cognitive assessment superior to the mini-mental state examination in detecting subtle cognitive impairment among middle-aged outpatient US Military veterans?. Archives of clinical neuropsychology, 27(7), 742–748.

129. Wolfsgruber, S., Jessen, F., Wiese, B., Stein, J., Bickel, H., Mösch, E., Weyerer, S., Werle, J., Pentzek, M., Fuchs, A. & Köhler, M. (2014). The CERAD neuropsychological assessment battery total score detects and predicts Alzheimer disease dementia with high diagnostic accuracy. The American Journal of Geriatric Psychiatry, 22(10), 1017–1028.

130. Wolpert, D. H., & Macready, W. G. (1997). No free lunch theorems for optimization. IEEE transactions on evolutionary computation, 1(1), 67–82.

131. Wright, A., Hickman, T. T. T., McEvoy, D., Aaron, S., Ai, A., Andersen, J. M., Hussain, S., Ramoni, R., Fiskio, J., Sittig, D.F. & Bates, D. W. (2016). Analysis of clinical decision support system malfunctions: a case series and survey. Journal of the American Medical Informatics Association, 23(6), 1068–1076.

