## Appendix 1 for "A practical computerized decision support system for predicting the severity of Alzheimer’s disease of an individual"

### **Appendix A**

**Table A.1.** Data types and their abbreviations.

| **Type** | **Feature** | **Abbreviation** | **Type** | **Feature** | | **Abbreviation** |
| --- | --- | --- | --- | --- | --- | --- |
| Socio-demographics & family history* | Age | Age | Neuropsychological/functional assessment | AD assessment scale-13 | | ADAS13 |
|  | Gender | PTGENDER |  | Mini-mental state examination | | MMSE |
|  | Education | PTEDUCAT |  | Functional activities questionnaire | | FAQ |
|  | Ethnicity | PTETHCAT |  | Montreal Cognitive Assessment | | MoCA |
|  | Race | PTRACCAT |  | Logical Memory Immediate Recall | | LIMMTOTAL |
|  | Marital status | PTMARRY |  | Logical Memory Delayed Recall | | LDELTOTAL |
|  | Dementia history from dad | FHQDAD |  | Rey auditory verbal learning test | Immediate | RAVLT Immediate |
|  | Dementia history from mom | FHQMOM |  |  |  |  |
|  | Does the participant have any siblings? | FHQSIB |  |  | Learning | RAVLT Learning |
| Medical history* | Psychiatric | MHPSYCH |  |  | Forgetting | RAVLT Forgetting |
|  | Neurologic | MH2NEURL |  |  | Percentage Forgetting | RAVLT Perc Forgetting |
|  | Head, eyes, ears, nose and throat | MH3HEAD | PET data | FDG PET | | FDG |
|  | Cardiovascular | MH4CARD |  | AV45 PET | | AV45 |
|  | Respiratory | MH5RESP |  | Left inferior temporal gyri | | Temporal Left |
|  | Hepatic | MH6HEPAT |  | Right inferior temporal gyri | | Temporal Right |
|  | Dermatologic-connective tissue | MH7DERM |  | Left angular gyri | | Angular Left |
|  | Musculoskeletal | MH8MUSCL |  | Right angular gyri | | Angular Right |
|  | Endocrine-metabolic | MH9ENDO |  | Bilateral posterior cingulate | | CingulumPost Bilateral |
|  | Gastrointestinal | MH10GAST |  |  |  |  |
|  | Hematopoietic-lymphatic | MH11HEMA |  | Sum of z-scores more than 2 standard deviations below the mean of normal control subjects | | SUMZ2 |
|  | Renal-genitourinary | MH12RENA |  | Sum of z-scores more than 3 standard deviations below the mean of normal control subjects | | SUMZ3 |
|  | Allergies or drug sensitivities | MH13ALLE | MRI data | Ventricles volume | | Ventricles |
|  | Alcohol abuse | MH14ALCH |  | Hippocampus volume | | Hippocampus |
|  | Drug abuse | MH15DRUG |  | Whole Brain volume | | Whole Brain |
|  | Smoking | MH16SMOK |  | Entorhinal volume | | Entorhinal |
|  | Malignancy | MH17MALI |  | Fusiform volume | | Fusiform |
|  | Major surgical procedures | MH18SURG |  | Middle temporal gyrus volume | | MidTemp |
|  | Other (if none, select ‘No’) | MH19OTHR |  | Intracerebral volume | | ICV |
| CSF biomarkers | Total tau protein (t-tau) | TAU |  | BSI whole brain volume | | BRAINVOL |
|  | Amyloid-β peptide of 42 amino acids (Aβ_1–42_) | ABETA |  | BSI ventricular volume | | VENTVOL |
|  |  |  |  | Cortical summary ROI (cortical grey matter regions of  frontal, anterior/posterior cingulate, lateral parietal, lateral temporal) divided by the whole cerebellum reference region | | WHOLECEREBNORM |
|  | Phosphorylated tau (p-tau_181p_) | PTAU |  | Cerebrospinal fluid volume | | CSF_V |
|  | Ratio of tau to Aβ_1–42_ | TAU_ABETA |  | Intracranial gray matter volume | | GRAY |
|  | Ratio of p-tau_181p_ to Aβ_1–42_ | PTAU_ABETA |  | Intracranial white matter volume | | WHITE |
|  |  |  |  | White matter hyperintensities (WMH) volume | | WHITMATHYP |
| * Medical and family history is either Yes or No. | | | | | | |

**Table A.2.** SVR and kNN_reg_ model performance measures for MRI, PET, CSF and cognitive function modalities retained for the training after feature selection. CFA represents 9 selected cognitive and functional assessments (LDELTOTAL, FAQ, MOCA, ADAS13, LIMMTOTAL, MMSE, RAVLT Immediate, RAVLT Perc Forgetting, RAVLT Learning), MRI - 4 features (Hippocampus, MidTemp, Entorhinal, Whole Brain), PET – 5 features (FDG, Angular Left, Angular Right, Temporal Left, SUMZ3), and CSF – 2 features (TAU_ABETA, TAU). ‘All’ features refer to a combination of MRI, PET, CSF, CFA, and Age. Performances of predictive models for each combination of modalities were recorded using an (unseen) testing set partitioned from the original data (10% of the original data). *R^2^*: adjusted coefficient of determination; RMSE: Root Mean Square Error. Asterix (*): a subset of features with the highest *R^2^*. For more details on data types and their abbreviations, refer to Table A.1.

| Features | SVR | | kNN_reg_ | |
| --- | --- | --- | --- | --- |
|  | *R^2^* | RMSE | *R^2^* | RMSE |
| All | 0.723, 95%CI (0.649,0.796) | 1.116 | 0.658, 95%CI (0.571,0.744) | 1.223 |
| CFA, PET, MRI, CSF | 0.736, 95%CI (0.662,0.809) | 1.082 | 0.641, 95%CI (0.548,0.734) | 1.274 |
| CFA, PET, MRI | 0.747, 95%CI (0.671,0.823) | 1.074 | 0.684, 95%CI (0.593,0.774) | 1.208 |
| CFA, PET, CSF | 0.789, 95%CI (0.720,0.858) | 0.975 | 0.637, 95%CI (0.530,0.744) | 1.277 |
| CFA, MRI, CSF | 0.788, 95%CI (0.716,0.859) | 1.010 | 0.723, 95%CI (0.633,0.812) | 1.103 |
| PET, MRI, CSF | 0.577, 95%CI (0.440,0.714) | 1.428 | 0.348, 95%CI (0.184,0.512) | 1.842 |
| CFA, PET | 0.788, 95%CI (0.714,0.862) | 0.985 | 0.662, 95%CI (0.555,0.770) | 1.228 |
| **CFA, MRI*** | **0.790, 95%CI (0.715,0.866)** | **1.004** | 0.738, 95%CI (0.647,0.829) | 1.074 |
| CFA, CSF | 0.767, 95%CI (0.680,0.854) | 1.040 | 0.708, 95%CI (0.604,0.813) | 1.131 |
| PET, MRI | 0.436, 95%CI (0.268,0.603) | 1.706 | 0.306, 95%CI (0.133,0.478) | 1.920 |
| PET, CSF | 0.257, 95%CI (0.079,0.435) | 1.946 | 0.361, 95%CI (0.180,0.542) | 1.703 |
| MRI, CSF | 0.525, 95%CI (0.359,0.692) | 1.537 | 0.480, 95%CI (0.306,0.654) | 1.132 |
| **CFA*** | 0.743, 95%CI (0.644,0.843) | 1.059 | **0.750, 95%CI (0.653,0.847)** | **1.043** |
| PET | 0.377, 95%CI (0.188,0.566) | 1.720 | 0.328, 95%CI (0.137,0.518) | 1.780 |
| MRI | 0.475, 95%CI (0.292,0.658) | 1.618 | 0.323, 95%CI (0.129,0.518) | 1.788 |
| CSF | 0.235, 95%CI (0.039,0.431) | 1.962 | 0.093, 95%CI (0.000,0.239) | 2.219 |
| Age | 0.139, 95%CI (0.000,0.077) | 2.148 | 0.013, 95%CI (0.000,0.074) | 2.156 |

**Table A.3.** RF model performance measures for MRI, PET, CSF and cognitive function modalities retained for the training after feature selection. CFA represents 9 selected cognitive and functional assessments (LDELTOTAL, FAQ, MOCA, ADAS13, LIMMTOTAL, RAVLT Immediate, MMSE, RAVLT Perc Forgetting, RAVLT Learning), MRI – 4 features (Hippocampus, MidTemp, Entorhinal, Whole Brain), PET – 5 features (FDG, Angular Left, Angular Right, Temporal Left, SUMZ3), and CSF – 2 features (TAU_ABETA, TAU). ‘All’ features refer to a combination of MRI, PET, CSF, CFA, and Age. Performances of predictive models for each combination of modalities were recorded using an (unseen) testing set partitioned from the original data (10% of the original data). MCA: multi-class classification accuracy. Multi-class AUC: multiclass area under the curve. Asterix (*): a subset of features with the best predictive performance. For more details on data types and their abbreviations, refer to Table A.1.

| Features | MCA (%) | Sensitivity (%) | | | | | Specificity (%) | | | | | | Multi-class AUC  (%) | |
| --- | --- | --- | --- | --- | --- | --- | --- | --- | --- | --- | --- | --- | --- | --- |
|  |  | Normal | | QCI | Mild/ Moderate | | Normal | | QCI | | Mild/ Moderate | |  |  |
| All | 78.7, 95%CI (64.3,89.3) | 82.4 | 80.8 | | | 50.0 | | 83.3 | | 76.2 | | 100 | | 85.7 |
| CFA, PET, MRI, CSF | 74.5, 95%CI (59.7,86.1) | 88.2 | 69.2 | | | 50.0 | | 73.3 | | 81.0 | | 100 | | 86.1 |
| CFA, PET, MRI | 78.7, 95%CI (64.3,89.3) | 82.4 | 80.8 | | | 50.0 | | 83.3 | | 76.2 | | 100 | | 85.7 |
| CFA, PET, CSF | 72.3, 95%CI (57.4,84.4) | 82.4 | 69.2 | | | 50.0 | | 73.3 | | 76.2 | | 100 | | 84.7 |
| CFA, MRI, CSF | 72.3, 95%CI (57.4,84.4) | 82.4 | 69.2 | | | 50.0 | | 73.3 | | 76.2 | | 100 | | 84.7 |
| PET, MRI, CSF | 53.2, 95%CI (38.1,67.9) | 76.5 | 42.3 | | | 25.0 | | 50.0 | | 66.7 | | 100 | | 78.2 |
| CFA, PET | 76.6, 95%CI (62.0,87.7) | 88.2 | 73.1 | | | 50.0 | | 76.7 | | 81.0 | | 100 | | 86.5 |
| CFA, MRI | 76.6, 95%CI (62.0,87.7) | 82.4 | 76.9 | | | 50.0 | | 80.0 | | 76.2 | | 100 | | 85.3 |
| CFA, CSF | 76.6, 95%CI (62.0,87.7) | 82.4 | 76.9 | | | 50.0 | | 80.0 | | 76.2 | | 100 | | 85.3 |
| PET, MRI | 44.7, 95%CI (30.2,59.9) | 82.4 | 26.9 | | | 0.0 | | 40.0 | | 66.7 | | 97.7 | | 76.9 |
| PET, CSF | 53.2, 95%CI (38.1,67.9) | 82.4 | 38.5 | | | 25.0 | | 46.7 | | 71.4 | | 100 | | 79.8 |
| MRI, CSF | 46.8, 95%CI (32.1,61.9) | 70.6 | 34.6 | | | 25.0 | | 53.3 | | 61.9 | | 93.0 | | 75.3 |
| **CFA*** | **80.0, 95%CI (66.7,90.9)** | **88.2** | **80.8** | | | **50** | | **83.3** | | **81.0** | | **100** | | **87.1** |
| PET | 51.1, 95%CI (36.1,65.9) | 82.4 | 34.6 | | | 25.0 | | 46.7 | | 71.4 | | 97.7 | | 79.3 |
| MRI | 48.9 95%CI (34.1,63.9) | 70.6 | 38.5 | | | 25 | | 50.0 | | 66.7 | | 95.3 | | 67.8 |
| CSF | 42.6, 95%CI (28.3,57.8) | 64.7 | 19.2 | | | 100 | | 63.3 | | 85.7 | | 69.8 | | 78.3 |
| Age | 38.3, 95%CI (24.5,53.6) | 52.9 | 30.8 | | | 25 | | 60.0 | | 66.7 | | 76.7 | | 58.5 |

**Table A.4.** kNN_class_ model performance measures for MRI, PET, CSF and cognitive function modalities retained for the training after feature selection. CFA represents 9 selected cognitive and functional assessments (LDELTOTAL, FAQ, MOCA, ADAS13, LIMMTOTAL, RAVLT Immediate, MMSE, RAVLT Perc Forgetting, RAVLT Learning), MRI – 4 features (Hippocampus, MidTemp, Entorhinal, Whole Brain), PET – 5 features (FDG, Angular Left, Angular Right, Temporal Left, SUMZ3), and CSF – 2 features (TAU_ABETA, TAU). ‘All’ features refer to a combination of MRI, PET, CSF, CFA, and Age. Performances of predictive models for each combination of modalities were recorded using an (unseen) testing set partitioned from the original data (10% of the original data). MCA: multi-class classification accuracy. Multi-class AUC: multiclass area under the curve. Asterix (*): a subset of features with the best predictive performance. For more details on data types and their abbreviations, refer to Table A.1.

| Features | MCA (%) | Sensitivity (%) | | | | | Specificity (%) | | | | | | Multi-class AUC  (%) | |
| --- | --- | --- | --- | --- | --- | --- | --- | --- | --- | --- | --- | --- | --- | --- |
|  |  | Normal | | QCI | Mild/ Moderate | | Normal | | QCI | | Mild/ Moderate | |  |  |
| All | 70.2, 95%CI (55.1, 82.7) | 94.1 | 53.9 | | | 75.0 | | 73.3 | | 90.5 | | 90.7 | | 88.4 |
| CFA, PET, MRI, CSF | 63.8, 95%CI (48.5,77.3) | 82.4 | 50.0 | | | 75.0 | | 70.0 | | 81.0 | | 90.7 | | 85.7 |
| CFA, PET, MRI | 78.7, 95%CI (64.3,89.3) | 94.1 | 65.4 | | | 100 | | 83.3 | | 95.2 | | 90.7 | | 93.4 |
| CFA, PET, CSF | 66.0, 95%CI (50.7,79.1) | 64.7 | 61.5 | | | 100 | | 73.3 | | 71.4 | | 95.3 | | 88.2 |
| **CFA, MRI, CSF*** | **89.7, 95%CI (76.9,96.5)** | **82.4** | **92.3** | | | **100** | | **96.7** | | **85.7** | | **97.7** | | **95.9** |
| PET, MRI, CSF | 57.5, 95%CI (42.2,71.7) | 76.5 | 42.3 | | | 75.0 | | 56.7 | | 85.7 | | 90.7 | | 74.6 |
| CFA, PET | 61.7, 95%CI (46.4,75.5) | 64.7 | 53.9 | | | 100 | | 73.3 | | 71.4 | | 90.7 | | 87.3 |
| CFA, MRI | 89.4, 95%CI (76.9,96.5) | 94.1 | 84.6 | | | 100 | | 93.3 | | 95.2 | | 95.3 | | 96.5 |
| CFA, CSF | 72.3, 95%CI (57.4,84.4) | 70.6 | 69.2 | | | 100 | | 80.0 | | 76.2 | | 95.3 | | 90.4 |
| PET, MRI | 63.8, 95%CI (48.5,77.3) | 82.4 | 53.9 | | | 50.0 | | 70.0 | | 85.7 | | 88.4 | | 73.5 |
| PET, CSF | 48.9, 95%CI (34.1,63.9) | 70.6 | 34.6 | | | 50.0 | | 53.3 | | 76.2 | | 88.4 | | 69.4 |
| MRI, CSF | 59.6, 95%CI (44.3,73.6) | 58.8 | 57.7 | | | 75.0 | | 70.0 | | 81.0 | | 86.1 | | 70.2 |
| CFA | 76.6, 95%CI (62.0,87.7) | 82.4 | 69.2 | | | 100 | | 80.0 | | 85.7 | | 95.4 | | 92.2 |
| PET | 42.6, 95%CI (28.7,57.8) | 58.8 | 30.8 | | | 50.0 | | 56.7 | | 61.9 | | 86.1 | | 73.9 |
| MRI | 55.3, 95%CI (40.1,69.8) | 64.7 | 46.2 | | | 75.0 | | 73.3 | | 81.0 | | 79.1 | | 72.1 |
| CSF | 29.8, 95%CI (17.3,44.9) | 41.2 | 19.2 | | | 50.0 | | 63.3 | | 61.9 | | 67.4 | | 60.3 |
| Age | 53.2, 95%CI (38.1,67.9) | 47.1 | 65.4 | | | 0.0 | | 80.0 | | 61.9 | | 81.4 | | 50.7 |
